# Quantifying electro-osmotic flow in mutant MspA nanopores

**DOI:** 10.64898/2026.11.10.765596

**Authors:** Tor Eriksson, Chenyu Wen, Cees Dekker

**Affiliations:** Department of Bionanoscience, Kavli Institute of Nanoscience Delft, Delft University of Technology, Van der Maasweg 9, 2629 HZ Delft, The Netherlands; Department of Electrical Engineering; Solid State Electronics, Uppsala University, 75121 Uppsala, Sweden

## Abstract

Nanopores have emerged as a powerful tool for single-molecule sensing and sequencing. In a typical nanopore experiment, a charged analyte such as DNA is driven through the pore by an electrophoretic force generated by the applied electric field. This can be problematic, however, when analytes such as proteins carry mixed charges. An alternative driving force can be from electro-osmotic flow (EOF), the water flow that arises from an asymmetric migration of cations and anions through the nanopore. However, despite much recent interest, there is no straightforward experimental method to quantify EOF in nanopores. Here, we trap a dipolar peptide in an MspA nanopore, where studying its escape rate versus voltage allows us to extract the work done by the EOF in various MspA mutants and wildtype MspA compared to a M2-MspA reference. For the 30 amino acid model peptide, this yields values of *∼*5.6-16.7 *k_B_T* for the work done by the EOF at 80 mV. Finite-element simulations confirm the trends observed in the experiments. Charged mutations close to the nanopore’s constriction site are found to produce the highest EOF. Our novel method provides a quantitative metric of the EOF in nanopores, opening up its study and broad use in future nanopore sequencing and sensing applications.

## 1 Introduction

Nanopores have emerged as a powerful tool for single-molecule sensing and DNA sequencing.[28, 56] In a typical nanopore experiment, a biological nanopore sits in a lipid bilayer separating two reservoirs filled with an electrolyte. When an electric field is applied across the membrane, ions flow through the nanopore, resulting in an ionic current (referred to as the open state current). The electrophoretic (EP) force can capture a molecular analyte such as a DNA molecule, and as the molecule traverses the nanopore it blocks some of the nanopore channel’s volume, resulting in a temporary current drop. By studying the features, such as depth, residence time, pattern, and noise fluctuations of this current drop, nanopores can distinguish fine molecular details differing by only a few atoms, providing the basis for sequencing.[46, 43, 30] Among the commonly used nanopores, the Mycobacterial Porin MspA is known for its well-defined constriction with a narrow (1.2 nm) diameter.[25]

While DNA nanopore sequencing has been achieved[35, 17], much research is now focused on developing single-molecule nanopore sequencing of proteins.[11, 41, 55, 14] Unlike DNA, however, proteins carry mixed charge and therefore cannot be uniformly captured and translocated by the EP force,[11, 41, 55] as that force will pull parts of the linearized protein in the opposite direction of the intended translocation. This may cause the peptide to become entangled above the pore, or to suddenly translocate in jumps, convoluting the signal.[16] In addition, the charge (as well as the hydrophobicity) of the analyte can induce undesired sequence-specific interactions to nanopore’s surface[11, 15] or intra-anaylte interactions. It is therefore of great interest to find new ways to more generally capture and translocate any analyte. For this purpose electro-osmotic flow (EOF) has gained much attention in recent years.[27, 40, 5, 29, 53, 49, 33, 42, 57, 47, 48] EOF is the water flow that arises due to an asymmetric migration of cations and anions in a solution. If, for example, a nanopore is anion-selective due to the positive surfaces charges on its inner lumen, more negative counter ions (i.e., anions) will be present in the pore. Under an applied electric field, these ions will flow through the pore and drag the surrounding body of water along with them via their hydration shells, and, accordingly, any analyte molecules will also be pulled along by the viscous shear force. If this EOF force exceeds the EP force, analytes could traverse the pore irrespective of their charge. Recently, EOF has been studied through simulations and experiments with ion-selective nanopores[47, 6, 42, 33, 9], guanidinium ions[57, 38], asymmetric or different salt conditions[13, 10, 39], and nanopore geometries[22].

However, much of this work remained qualitative as there is no method to directly quantify the strength of the EOF on the analyte experimentally. Computational methods such as full-atom molecular dynamics may provide estimates of the EOF, but it remains uncertain how well they mimic reality.[38, 6] On the experimental side, measurements of EOF have been qualitative, *i.e.* addressing whether or not a certain charged peptide can translocate the nanopore against an opposing EP force[47], or using proxies such as the capture rate of an analyte.[13] Some measure of the EOF strength can be obtained from the ion selectivity of a nanopore, which is estimated from the reversal potential generated using a salt gradient across a nanopore. This, however, comes with some limitations. First, the ion selectivity is still not more than an indirect rough proxy for the EOF, as recently discussed in depth by Di Muccio *et al*.[21] Second, while the ion selectivity can distinguish differences between pores, it does not capture how the EOF behaves at various voltages or buffer conditions. In addition, it is uncertain how EOF estimated by the ion selectivity translates to the force on an analyte in a nanopore. Summing up, there is a need to develop a reliable metric to quantify EOF.

Here, we develop such an EOF metric by applying ‘nanopore tweezers’, in which a dipolar peptide is trapped in the nanopore due to opposing EP forces at the two ends of the peptide.[18, 4, 45] Nanopore tweezers were originally developed to trap a molecule of interest within a nanopore, enabling prolonged study of the molecule. Here, we use it in another way, as we show that we can quantify the EOF by measuring the escape rate of the analyte from the trap (Fig.1). We measured the trapping times of the dipolar peptide for several MspA mutants, that generate EOF with different strengths, and compared those to M2-MspA as a reference. This allowed us to quantify the work done by EOF on the analyte in five different MspA mutants as well as in wildtype (WT) MspA, yielding values for the work done by EOF on the peptide of *∼*5.6-16.7 *k_B_T* at 80 mV. We thus establish, to our knowledge, the first widely applicable experimental method to quantify the work performed on a peptide by EOF in a nanopore.

**Figure 1:**
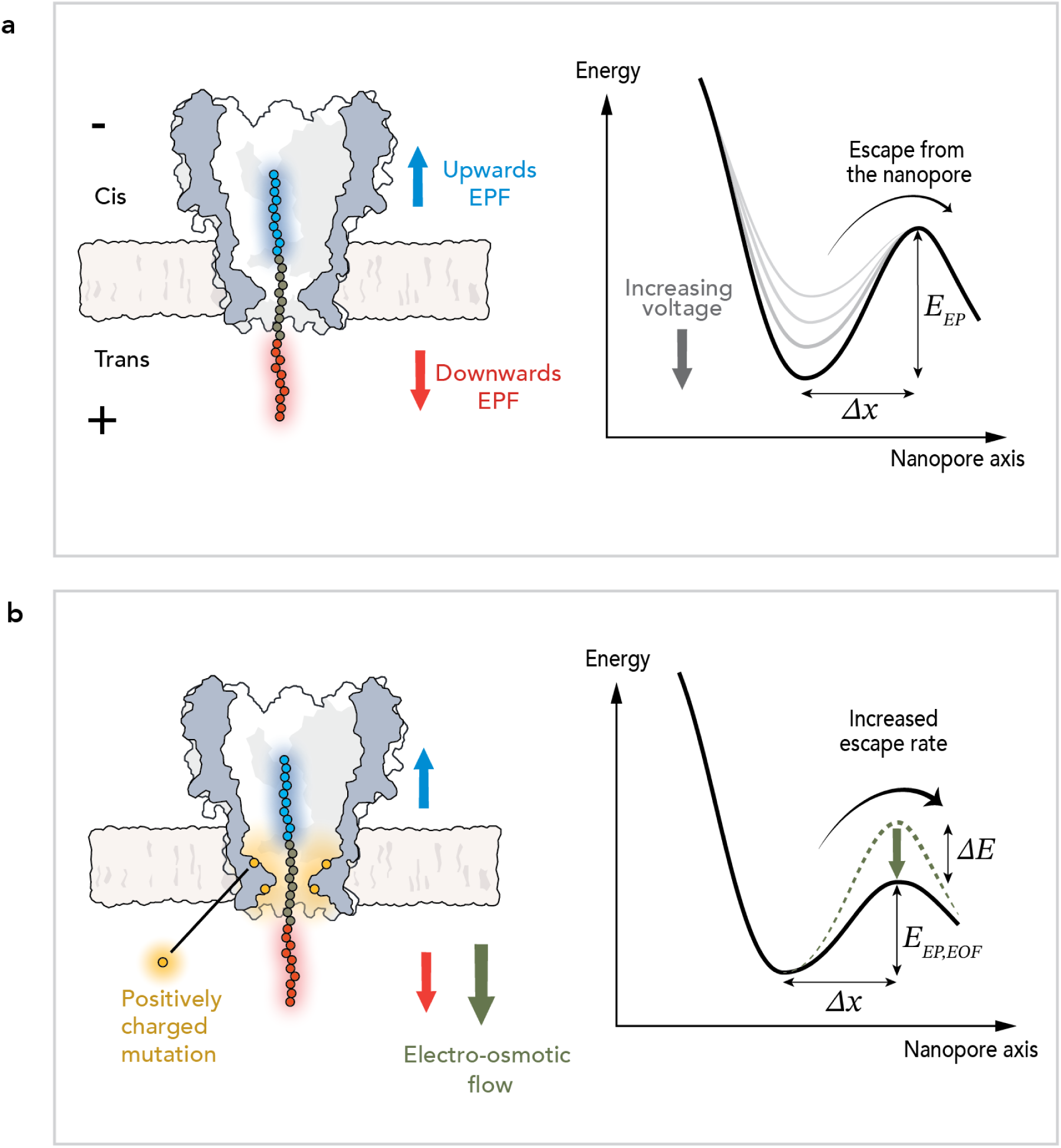
Overview of the nanopore tweezer system. a) In a nanopore tweezer system, a dipolar peptide is trapped between opposing electrophoretic forces (*F_EF_*) (left). The escape of the peptide from the nanopore can be modeled as a energy-barrier-crossing process (right), where the energy well deepens at higher electric fields. b) By adding electrosmotic flow (EOF) to the system, for example by introducing charged mutations in the nanopore lumen, the peptide will feel an additional unidirectional downward force from the EOF (left). The additional force from the EOF modulates the energy barrier, altering the escape rate (right).

We also compare these findings with the ion selectivity measured for these nanopores and find that ion selectivity has a monotonic but strongly nonlinear relationship to the actual work performed by the EOF. The findings provide a quantitative metric for the EOF in nanopores, opening up its study and broad use in future nanopore sequencing and sensing applications.

## 2 Results and Discussion

### 2.1 Modeling EOF on a dipolar peptide in a nanopore

We realized that ‘nanopore tweezers’ measurements can yield a quantitative measure of the strength of the EOF in nanopores. In these tweezers, a dipolar peptide such as E_10_N_10_R_10_ (*i.e.*, a string of 30 amino acids of negatively charged glutamic acid (glutamate; E), neutral asparagine (N), and positively charged arginine (R)) is trapped in a nanopore due to opposing EP forces, see Fig. 1a. In passing, we note that this differs from the picometer-resolution nanopore force-spectroscopy approach where the analyte is trapped between the constriction and a motor enzyme - which was also named ‘nanopore tweezers’[20]. One force pulls the negatively charged bottom end of the peptide downward, while another force pulls the positively charged end upward, see Fig. 1a, left.[18, 45, 4] The peptide is thus trapped at the nanopore’s constriction site because the EP forces balance each other (Fig. 1a). Because the peptide is stretched under opposing EP forces, it mimics the peptide sequencing system[11, 55] quite well, in which the analyte is similarly stretched between a downward EP force and an upward pulling force from a motor enzyme.

We can model the trapping and escape of the peptide as an energy-barrier-crossing process, in which the peptide must overcome an energy barrier *E* to escape from the nanopore at a rate *k*. The electric field is strongest at the constriction site, and hence the peptide is trapped at that position in an energy well. For stronger applied electric fields, the EP forces that hold the peptide are also stronger, yielding a higher energy barrier and a lower escape rate (Fig. 1a, right). Assuming that the peptide only escapes from one side of the nanopore (which we show to be true for MspA, see below), we can describe this as a single-rate escape process[8, 37, 23, 24, 53]

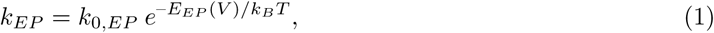

where *k* is the measured escape rate, *k*_0*,EP*_ is the escape rate at *V* = 0, and *E_EP_*(*V*) is the energy barrier height which depends on the applied voltage.

If there is a substantial EOF in the nanopore, the peptide will experience an additional downward force (Fig. 1b, left), and hence the escape rate can be described by

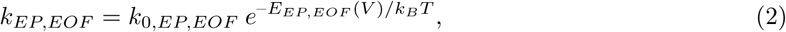

where the quantities are analogous to those of Eq. 1 but for a condition with EOF. As the EOF force is directed downward, it lowers the barrier for escape (Fig. 1b, right). The energy difference between the barriers in the two conditions is the *work W_EOF_* done by EOF on the peptide, i.e.,

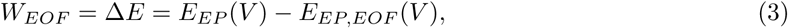

where Δ*E* is the difference in the energy barrier between the two conditions.

Four things are worth noting. First, as apparent from Eq. 3, our method does not extract the absolute amount of EOF work on the peptide, but the *additional* EOF work compared to a reference. Ideally, that reference is selected to have a negligible intrinsic amount of EOF. This appears to be the case for the M2-MspA variant[6, 32, 38], and as a result our approach is well suited to estimate the strength of the EOF for MspA mutants. Second, this approach assumes that the EP force is the same in the two conditions, which is a reasonable assumption, as a single pore mutation that causes EOF does not significantly change the pore’s geometry and the electric field inside the nanopore. Third, *k*_0_ describes the rate of escape at *V* = 0, which involves electrostatic interactions, hydrophobic interactions, and entropy. While *k*_0_ is independent of *W_EOF_*, it may be of interest for understanding these quantities. Fourth, our method does not quantify the EOF as a flow velocity, but as the work that is performed on the peptide by EOF. This can be described by

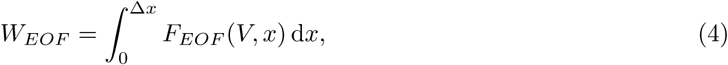

where *F_EOF_* (*V, x*) is the force applied to the peptide at voltage *V* at a distance *x* from the energy minimum, and Δ*x* is the physical distance between the energy minimum and the energy barrier for escape - see Fig. 1. Altering the peptides charge or length would change both the transition-state distance Δ*x* and the force profile *F_EOF_* (*V, x*), giving a different value for the work, and hence the same peptide must be used in both the reference and the measurement.

In summary, nanopore tweezers may be used as a tool for experimentally determining the work done by EOF, something that previously was not possible.

### 2.2 Dipolar peptides exit MspA from the *trans* side

We first investigated the peptide E_10_N_10_R_10_ (Fig. 2a) in the M2-MspA nanopore. We chose M2-MspA as our reference due to its low EOF.[6, 32, 38] To verify that the M2-MspA worked as a nanopore tweezer for the peptide, we measured the trapping times, and its inverse, the escape rates. All measurements in this study were done at 900 mM KCl, unless specified otherwise. Upon application of a 100 mV voltage, we saw clear, well-defined trappings - see Fig. 2b. The open state current of about 180 pA (*I*_0_), was observed to suddenly drop to 50 pA (*I*) when a peptide was trapped in the nanopore, whereupon the current level reversed to the open state current after some time (*∼*1 second) when the peptide escaped the trap. Upon decreasing the applied voltage from 120 mV to 80 mV, the trapping times decreased by approximately two orders of magnitude (Fig. 2c & d, & Fig. S4). The relative current blockade, *I/I*_0_, however, changed only slightly from 0.28 to 0.24. The particular peptide E_10_N_10_R_10_ was chosen because it showed a good capture rate and trapping times at 100 mV, which is an applied voltage that is commonly used in nanopore experiments.

**Figure 2:**
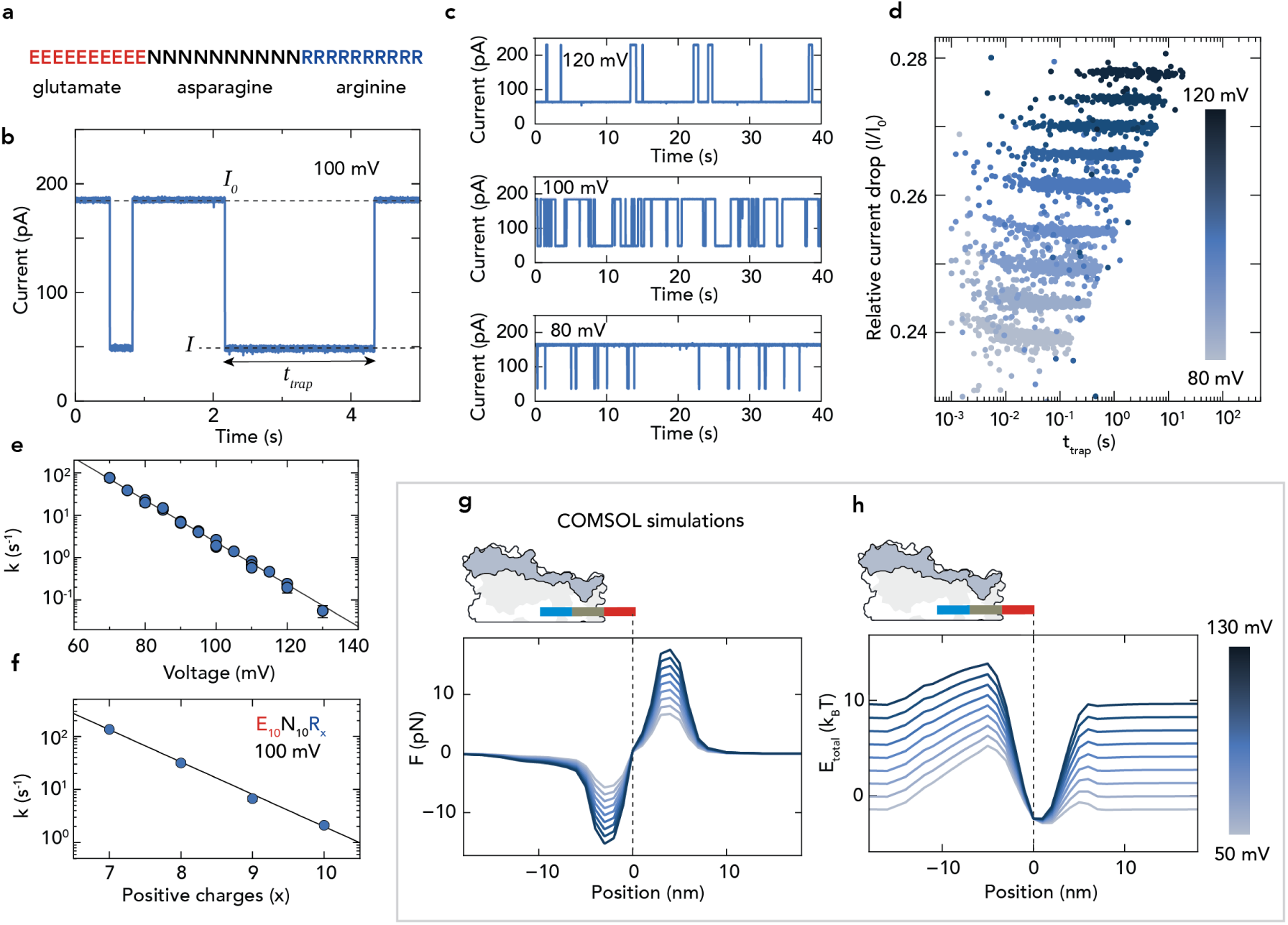
Nanopore tweezer data for the M2-MspA nanopore. a) Sequence of the bipolar peptide used in this study: a 30 amino-acid string consisting of 10 glutamate (E), 10 asparagine (N), and 10 arginine (R). b) Example current trace with two trapping events of the peptide in the M2-MspA nanopore at 100 mV. The open state current *I*_0_ of about 180 pA is seen to suddenly drop to an event current *I* of about 50 pA when the peptide is trapped, whereupon it returns to the open state current when the peptide escapes the nanopore. *t_trap_* denotes the time duration that the peptide is trapped in the nanopore. c) Example current traces at 120, 100, and 80 mV. By decreasing the voltage, the EP force trapping the peptide in the pore becomes weaker, thus decreasing the trapping times. d) Scatter plot of all trapping events in M2-MspA across different voltages for one replicate. With increasing voltage we see the trapping times become much longer and the relative event current become slightly larger. A histogram of the distribution at each voltage is provided in Fig. S4. e) Semilog plot of the mean escape rate vs. voltage, showing an exponential dependence. Standard error of the mean error bars are smaller than the markers when not visible. The line describes *ln*(*k*) = *ln*(*k*_0_) *− p*_0_*V*. f) Escape rate at 100 mV for the peptide E_10_N_10_R_x_, where x = 7, 8, 9, 10. For decreasing positive charge, the escape rate of the peptide increased, consistent with a decreasing EP force directed towards the *cis* side. Standard error of the mean error bars are smaller than the markers when not visible. g) The total force on the peptide was estimated using numerical simulations. The peptide was approximated as a stiff dipolar rod that moved along the nanopore axis in 1 nm steps. x-axis denotes position along the nanopore’s central axis, where 0 is the energy minimum. Positive and negative force values indicate that the net force is directed towards the *cis* and *trans* side, respectively. h) Same as g, but for the total energy landscape including entropy.

To see if the escape rates would fit the expected behavior described by Eq. 1, we measured the voltage dependence of the peptide escape rate. Experimentally, the lower voltage limit was set by the minimal voltage where peptide capture was possible, and the upper voltage limit was set by the occurrence of events where multiple peptides were trapped in the pore simultaneously (see supplementary section S1.1 for a detailed description of this phenomenon).

At each voltage, the distribution of trapping times was extracted by binning the trapping times in a histogram, and subsequently performing a least-squares fit of a single exponential decay - an approach that was used before in similar nanopore experiments.[34, 26]. This yielded the characteristic trapping time, and hence its inverse, the mean escape rate. When plotting the logarithm of the mean escape rate against voltage, a linear decrease was observed (Fig. 2e), i.e.

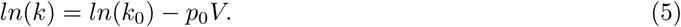

This linear fit with fitting parameters *k*_0_ and *p*_0_ described our data well (*χ*^2^ = 0.54, *p* = 0.81; see Tab.S1 for fitting parameters). The same relationship is reported in similar trapping experiments.[18, 4] Note that this single-barrier crossing model assumes that the peptides only exit the nanopore through the *trans* exit. To test this assumption, we measured the escape rates of a series of peptides E_10_N_10_R_x_, where x = 7, 8, 9, 10. With a decreasing number of positive charges, we observed an exponential increase in the escape rate (Fig. 2f). This indicates that the force keeping the peptide trapped in the nanopore decreased for lower *x* values. Because the EP force pulls the positive end of the peptide upwards towards the *cis* side, we concluded that the peptides exit through the opposing *trans* exit.

To verify that this unidirectional exit was a consequence of the MspA nanopore’s conical shape, we simulated the system using numerical simulations with COMSOL Multiphysics 6.1.[19] The peptide was modeled as a dipolarly charged rod that was translocated through the nanopore in 1 nm steps, with the forces and electrostatics simulated at each step (see the methods section for details). Fig. 2g depicts the force on the peptide. We observe a zero net force when the rod sits in the middle of the constriction site, consistent with the notion that the peptide is trapped at the constriction site of MspA, where the electric field is the strongest.

The energy barriers to the left and right deduced whether the escape was favored towards the *cis* or *trans* side, respectively. Fig. 2h depicts the energy landscape and shows that the energy barrier towards the *trans* side is lower, thus favoring a *trans*-side escape rather than a *cis*-side escape. We can in fact separate the energetic contributions from the EP forces and from entropy (Fig. S5a & b, respectively; see supplementary section S1.2 for the estimation of the entropy). We find that both of them independently favor an escape towards the *trans* side. Taken together, the data indicate an energy landscape that clearly favors *trans*-side escape of the peptides from MspA.

### 2.3 Extraction of the EOF work in the S103K-MspA mutant

Next, we tested whether we could measure the work done by EOF in the S103K-M2-MspA (hereafter referred to as S103K), an MspA mutant that is expected to exhibit strong EOF. The S103K mutation is set in the M2-MspA background. Because this mutant carries an additional positively charged residue located below the constriction site in each monomer, it exhibits EOF, as has been reported previously (Fig. 3a).[6]

**Figure 3:**
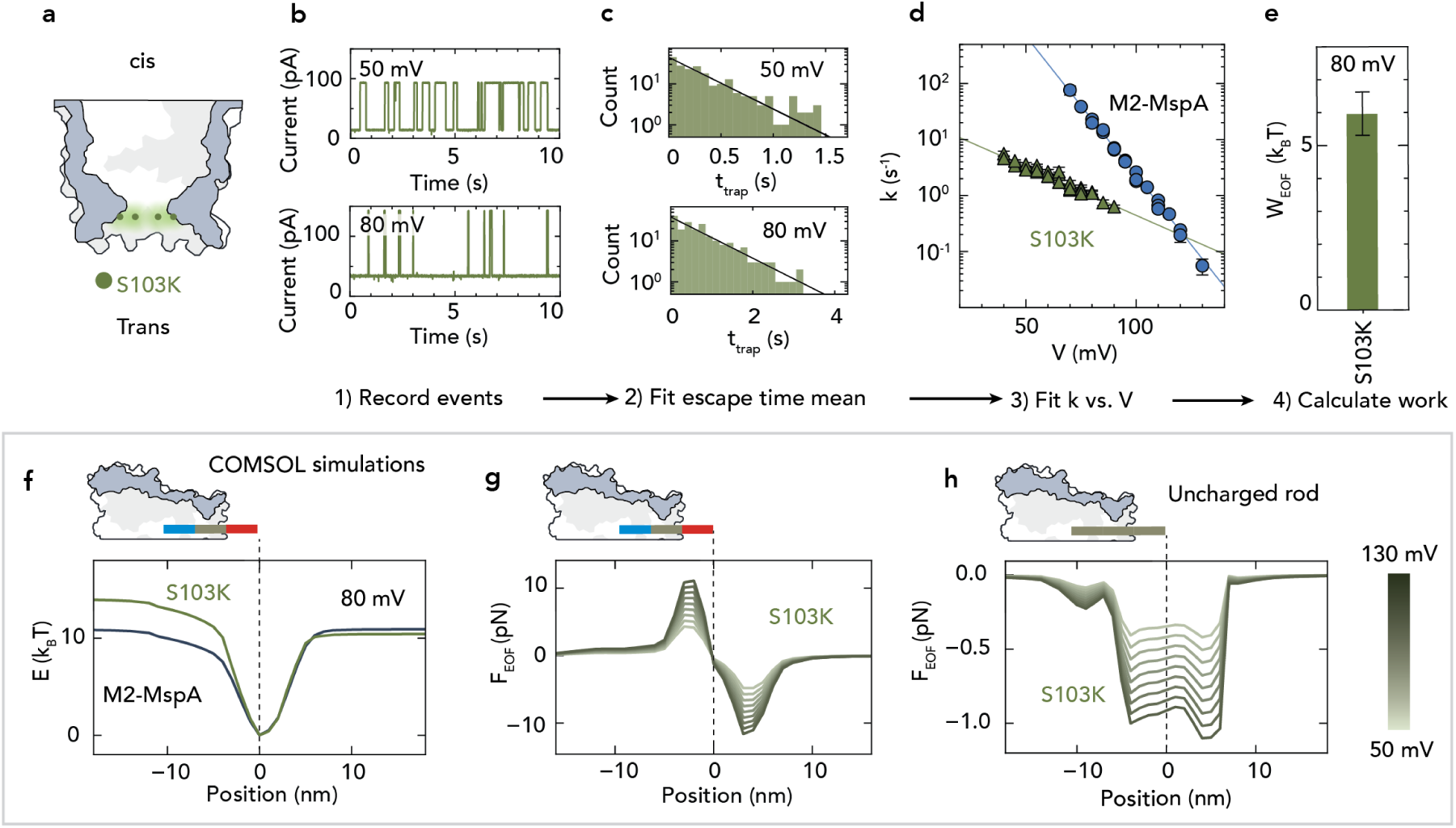
Extraction of work done by EOF in the S103K MspA mutant. a) Cartoon showing the position of the charged amino acids in the S103K mutant in the M2-MspA background. Panels b-e describes the workflow in extracting the work done by EOF. b) Example current traces at 50 and 80 mV. Trapping times were longer at 80 mV than at 50 mV. c) Example histograms of the current distribution at 50 and 80 mV. For each voltage the currents were binned into a histogram and the mean and standard error of the mean extracted by a *χ*^2^ fit of a single exponential decay process described by the black line. d) Escape rate vs. voltage for the S103K mutant and the M2-MspA. The escape rate in the S103K mutant decreased less with voltage, indicating the presence of EOF. The line describes *ln*(*k*) = *ln*(*k*_0_) *− p*_0_*V*. e) Bar graph of the work done by EOF in the S103K mutant. The work was calculated by taking the difference in how the energy barrier changes for the S103K mutant compared to the M2-MspA nanopore which has neglible EOF, i.e. *W_EOF_* = *k_B_T* (*p*_0_*_,M_*_2_ *− p*_0_*_,mutant_*)*V*. f) Energy landscape for the S103K mutant estimated using numerical simulations. y-axis denotes the energy in of the system excluding entropy; the x-axis denotes the rod position along the nanopore axis, with 0 being the energy minimum. g) Same as f, but the for EOF force in the S103K mutant. Positive and negative values indicate that the net force is directed towards the *cis* and *trans* side, respectively. The two peaks stem from the EOF induced by charges on the bipolar peptide. h) Same as g, but for an uncharged rod, allowing estimation of the EOF induced by the S103K mutant pore alone, without the influence of the peptides own charge.

The workflow for quantifying the EOF can be seen in Fig. 3b-e, where steps 1-3 (Fig. 3b-d) are analogous to those for the M2-MspA in the previous section. Fig. 3b shows typical current traces of the trapped peptide at 50 and 80 mV. We clearly see that increasing the voltage prolonged the trapping times. The upper voltage limit of 90 mV was here set by the trapping becoming to long for efficient capture. Fig. 3c shows the distribution of the trapping times for the extraction of the mean trapping time. Importantly, we observe that the slope in Fig. 3d for the S103K mutant data is less negative compared to the slope of the M2-MspA data (Tab.S1). A less negative slope means that the escape rate decreases slower with respect to voltage. This makes sense because the EOF force pushes the peptide out of the *trans* side, which is antagonistic to the EP force that is keeping the peptide trapped. It is also the key point, which makes it possible for us to combine Eq.(1)-(3) and (5) to calculate the work done by EOF as

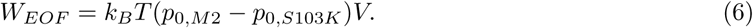

Note that this is valid because the EOF in the M2-MspA is near negligible. From the data in Fig. 3d we obtain a value of 6.0 *±* 0.7 *k_B_T* for the work done by EOF at 80 mV (Fig. 3e). Details about the experiments can be found in supplementary section S1.6.

Numerical simulations provide insights into the interplay of the EP force and the EOF force. Comparing the energy landscape between the S103K and M2-MspA nanopores (see Fig. 3f and Fig. S6a & b), the *trans*-side exit is even more pronounced for the S103K mutant because of the EOF. Numerical simulations allow us to examine the individual contributions from the EP and EOF. While the EP force, *F_EP_*, was expected to be very similar to that in the M2-MspA (Fig. S10c), we intuitively imagined that the profile of EOF force, *F_EOF_*, would appear as a negative plateau for the region where the peptide is located in the pore. Contrary to our expectation, however, *F_EOF_* showed a distinct structure along the vertical axis, displaying a negative and positive peak, in the reverse order to those for *F_EP_* (Fig. 3g & Fig. S6d). We attribute these peaks to the EOF induced by the charged ends of the peptide, as they pass through the strong electric field at the constriction site. The reason why the peaks appear in the opposite order to those for *F_EP_*is that a peptide segment of a particular charge will generate an EOF in the opposite direction to the EP force pulling on the segment. The variations in the EOF induced by the translocating peptide appeared to dominate the EOF induced by the nanopore. To obtain the force from the EOF generated by the nanopore itself, we simulated the same system but with an uncharged rod. As seen from Fig. 3h, we now obtained a more homogeneous negative profile of *F_EOF_*as expected. Because there is no electrophoretic tug-of-war on the uncharged peptide, the energy declines monotonically as the peptide traverses the pore *cis* to *trans* (Fig. S6e).

The numerical simulations thus indicate that the *F_EOF_* induced by the charged ends of the peptide (*F_EOF_∼*10pN) is much larger than the EOF generated by the nanopore (*F_EOF_ ∼*1pN). Note, however, that the same peptide is used in both the reference and the measurement, thereby canceling out the peptide-end contributions to the EOF. A limitation of the numerical simulations is also that they treat the ion distribution as a continuous field, an approximation that breaks down on the 1 nm scale of the nanopore. This leads the simulations to underestimate the effect of charges that are located further from the constriction site. Therefore, the *F_EOF_* from the S103K mutation is likely to be larger than that predicted by the numerical simulations (also see Fig. S10). The numerical simulations, however, do still provide qualitative insights that is meaningful to grasp the major physics.

### 2.4 Non-linear scaling of EOF in the S103K/N108K-MspA mutant

When we expanded our study to the S103K/N108K mutant of MspA, we observed a non-linear scaling of the EOF. The S103K/N108K double mutant was chosen to assess how an additional mutation influenced the EOF. In addition to the S103K mutation, it carries the N108K mutation which is located above the constriction site (Fig. 4a, inset). Thanks to the EOF-enhanced capture rate in this pore, we were able to record trappings and escapes for voltages as low as 25 mV, at which point the events became too fast to record. We expected the escape rate to depend linearly on voltage as it had in previous pores. However, the relationship turned out to be both nonlinear and to consist of two different regimes (Fig. 4a).

**Figure 4:**
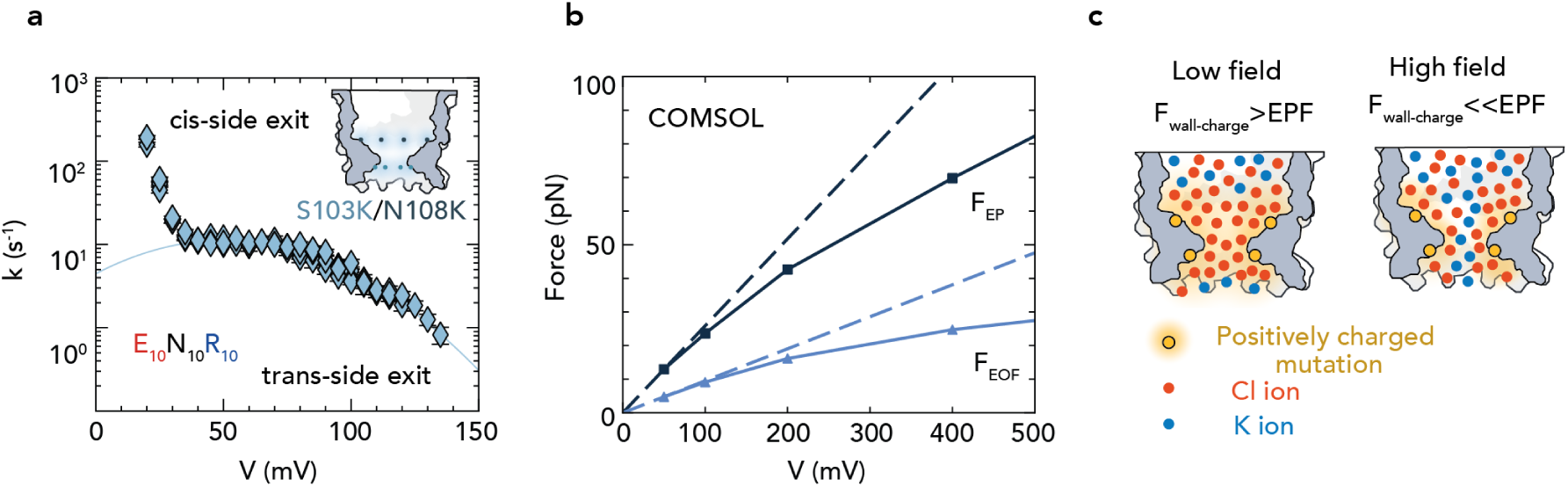
EOF scales nonlinearly with voltage in the S103K/N108K mutant. a) Escape rate vs. voltage for the S103K/N108K mutant. Inset shows the positions of the charged mutations set in the M2-MspA background. Below 40 mV the peptide exits out the *cis* side due to electrostatic repulsion between the peptide and the charges in the nanopore. Above 40 mV the peptide exists from the *trans* side of the pore due to the EP force and EOF. b) Numerical simulation of *F_EP_*and *F_EOF_*as the peptide passes through the pore. Both *F_EP_*and *F_EOF_*deviate from the linear trend above about 100 mV. c) Cartoon showing the influence of a strong electric field on the local ion concentration, as described by Li and Muthukumar[32] At low electric fields (left), the Coulomb force to the wall charge dominates, resulting in a high ion concentration localization. In a high electric field (right), the EP force dominates, resulting in weaker ion concentration polarization. This causes the effective ion selectivity to decrease and the EOF to weaken at higher fields.

Below 40 mV, we observed a sharp increase in the escape rate as the voltage was lowered. As both the EP force and EOF are weak at these low voltages, we conjectured that the electrostatic repulsion dominated here, where positive charges within the pore lumen repelled the peptides positive end, expelling it back into the *cis* side. This was not observed for the previous pores, as this positive charge is absent in the M2-MspA, and the S103K mutant was only measured down to 40 mV. For voltages above 40 mV, the EP and EOF forces dominated, and the peptide exited from the *trans* side. Unlike the S103K mutant, in this regime the S103K/N108K mutant showed a nonlinear dependence on voltage, with the slope becoming more negative at higher voltages.

In fitting the escape rate for the *trans*-side exit, we therefore added a non-linear term,

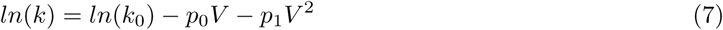

where *k*_0_, *p*_0_, and *p*_1_ are fit parameters. To only fit the *trans*-side-exit regime we fitted voltages equal or larger than 40 mV (Tab.S2).

To identify the source of this nonlinearity, we obtained the scaling of *F_EP_* and *F_EOF_* between 50 and 1000 mV using numerical simulations. Interestingly, we observe that both *F_EP_* and *F_EOF_* displayed a nonlinear dependence on the applied voltage beyond about 100 mV (Fig. 4b), with *F_EOF_*diverging more strongly than *F_EP_*. This is reminiscent of previous simulations by Li and Muthukumar.[32] Using a continuum model, they showed even stronger deviations, with the EOF velocity returning even to net 0 before 1000 mV in some MspA variants. They suggested that the nonlinearity arises from the interplay between the EP force and the wall charge: at low voltages, the electrostatic repulsion to the wall charge dominates the EP force, which leads to charge localization and the ion selectivity that generates EOF (Fig. 4c, left panel). With increasing voltage, however, the EP force becomes stronger while the electrostatic forces remain unchanged. Consequently, the EP force will drag many counter ions through the nanopore (Fig. 4c, right panel), thus reducing the nanopores ion selectivity and yielding a smaller EOF. Willems *et al.*, also using a continuum model, similarly observed how the local anion concentration is more sensitive to voltage than the cation concentration for a cation selective pore.[54]

With our charged analyte in the pore, an additional electrostatic repulsion might contribute as well. As the stronger electric field reduces the extent of the ion concentration localization in the pore, the Debye screening length becomes longer. Accordingly, the Coulomb force between the pore’s charge and the analyte becomes larger, adding another contribution to the energy barrier. If this effect has a considerable contribution, our assay would slightly underestimate *W_EOF_*. However, as we will see in Fig. 5c, *W_EOF_* in the measured range only deviates slightly from a linear relationship. At high voltage in short nanochannels, a reverse flow might arise because of a pressure gradient as a direct result of the EOF. This might also contribute to the nonlinear effect and is covered in supplementary section S1.4.

**Figure 5:**
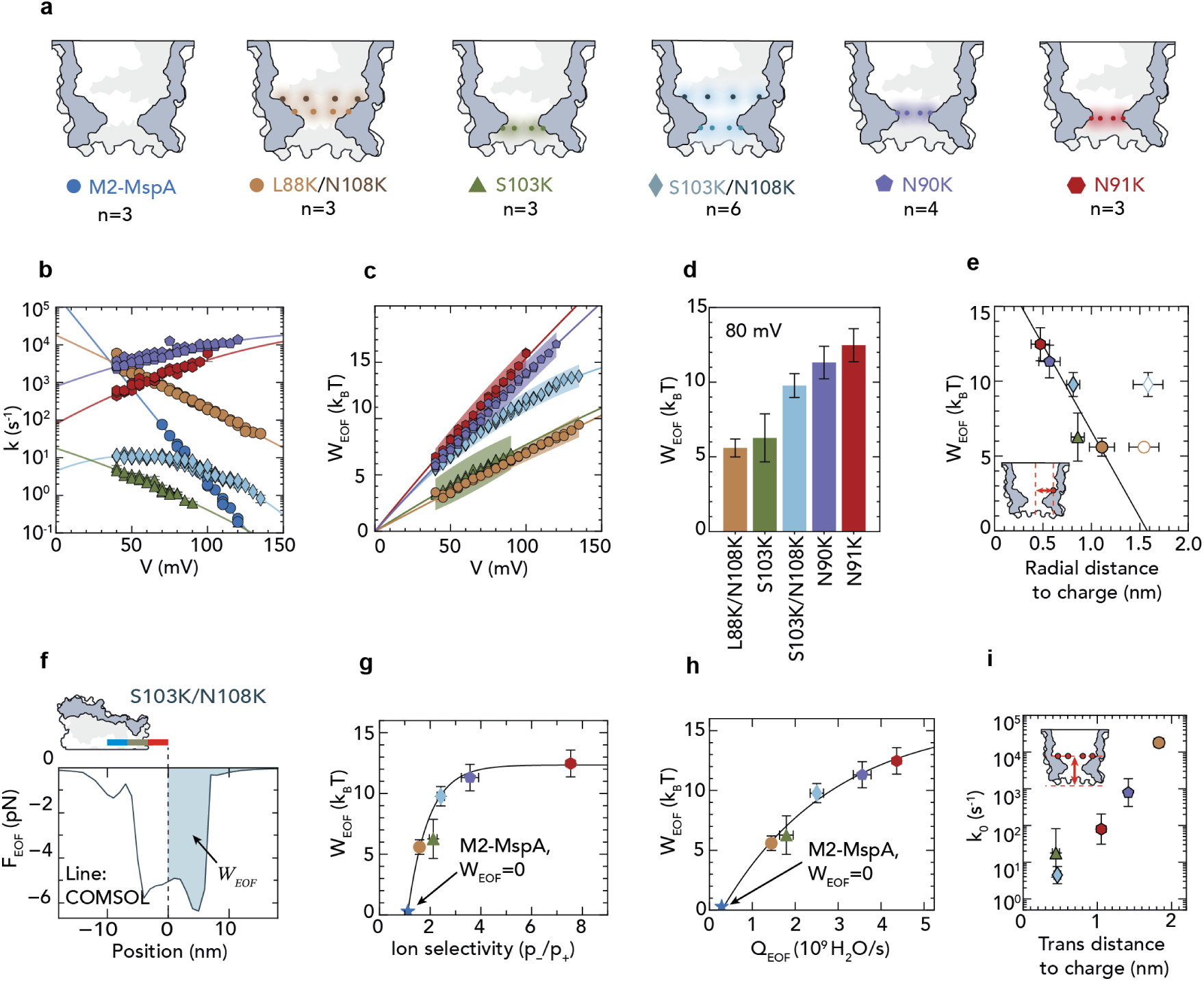
Comparison of work done by EOF in five different charged MspA mutants. a) Cartoon indicating the positions of the charges in the nanopores. The mutations are set in the background of the M2-MspA variant. n indicates the number of replicates in our experiments on these mutants. b) Escape rate vs. voltage. Only the values used for the fitting are shown (*trans*-side exit, *V ≥*40 mV). A plot of all data points can be found seen in Fig. S7b. MspA mutants are fit to *ln*(*k*) = *ln*(*k*_0_) *− p*_0_*V − p*_1_*V* ^2^ and M2-MspA is fit to *ln*(*k*) = *ln*(*k*_0_) *− p*_0_*V* (Tab.S1 & S2). Replicates were fitted individually and then a weighted average was calculated (see Fig. S7c). Error bars indicate standard error of the mean. c) Work done by the EOF for the MspA mutants across voltages. The work is obtained by taking the difference in how the energy barrier changes (Eq. 8), i.e. *W_EOF_*= *k_B_T* [(*p*_0_*_,M_*_2_ *− p*_0_*_,mutant_*)*V − p*_1_*_,mutant_V* ^2^]. The nonlinear effect discussed above only has a weak influence on *W_EOF_* below 150 mV. Shaded area indicates Standard deviation. d) Bar graph of the *W_EOF_* at 80 mV, calculated with the same formula as in panel c. Error bars indicate Standard deviation. e) Work done by EOF at 80 mV plotted against the radial distance of the charged atom closest to the nanopore axis. The line describes a linear fit (non-filled points not included). For the L88K/N108K and S103K/N108K double mutants the charges closest to the axis were used (L88K and S103K, respectively). The N108K mutation is plotted as an empty circle or diamond for the L88K/N108K and S103K/N108K mutants, respectively. Positions of the charged atoms were obtained using alpha fold 3[1] to predict the structures of the nanopores, and then taking the mean distance between the central axis to the charged atom in each monomer. Error bars indicate Standard deviation. f) Estimation of the force on the peptide along the central axis of the S103K/N108K mutant. The force profile was obtained from a numerical simulation by moving an uncharged rod along the S103K/N108K mutant in 1 nm steps. The area on the right of the dashed line (shaded area) corresponds to the *trans*-side exit was scaled to fit the experimentally measured work. This gave an estimate of the EOF force *F_EOF_* on the analyte of *∼*5 pN. The peak at approximately *∼*5 nm is an artifact from the EOF pushing on the blunt end of the rod when located in the constriction site. g) Work done by the EOF vs. the ion selectivity. The ion selectivity of the different pores was estimated by measuring the reversal potential in the nanopores over a 2 M KCl *cis*: 500 mM KCl *trans* salt gradient, and then using the GHK equation to calculate the ratio of the ion permeabilities. As the work done by the M2-MspA by definition is zero, it sits on the x-axis at the point of its ion selectivity (blue star). We observe a strongly nonlinear saturating trend between *W_EOF_* and the ion selectivity. Error bars indicate Standard deviation. The line is fitted to the saturating expression *W_EOF_* = *a ∗ e^−b∗Sp−/p^*^+^ + *c*, where *a*, *b*, and *c* are fit parameters, and *S_p−/p_*_+_ is the ion selectivity. h) Work done by the EOF vs. the water flux through the nanopores estimated from the ion selectivity by using the equation from Gu *et al.*[27] Again, a nonlinear relationship is observed. Like in panel g, The M2-MspA sits on the x-axis (blue star). Error bars indicate Standard deviation. The line is fitted to the same equation as in (g), but using the water flux *Q_EOF_* instead of the ion selectivity. i) *k*_0_, the escape rate at *V* = 0, vs. axial distance to the *trans* side. As the charges are progressively placed closer to the *trans* side of the nanopore, *k*_0_ decreases. This is due to the electrostatic repulsion pushing the peptide back into the nanopore. Distances to the *trans* side were obtained like in e, with the *trans* side defined as a plane orthogonal to the nanopore axis running through the lowest atom in the structure. For the L88K/N108K and S103K/N108K double mutants, the charges closest to the *trans* side are plotted (L88K and S103K, respectively). Error bars indicate Standard deviation.

In the investigation of the nonlinear behavior, we also measured the peptides E_10_N_10_R_x_ with x = 7, 8, 9 in S103K/N108K at 900 mM KCl, as well as the peptide E_10_N_10_R_8_ at various KCl concentrations - see supplementary sections S1.3 & S1.4, respectively.

### 2.5 Charges near the nanopore constriction site yield the highest EOF

Finally, we measured the L88K/N108K, N90K, and N91K MspA mutants, to compare a broad range of MspA mutants and to study the influence of the charge position on the EOF. The L88K/N108K mutant has both of its mutations above the constriction site, while the N90K and N91K mutations make up, respectively, the second narrowest and the narrowest parts of the constriction site itself (Fig5a).[6] All these three pores showed a rapid *cis*-side exit below 40 mV (Fig. S7b). We therefore focused the analysis on the *trans*-side escape regime above 40 mV and used values equal to or above this value for fitting. For consistency, we used the nonlinear expression (Eq. 7) for all mutant pores. The M2-MspA data still exhibited a linear dependence on voltage, and the nonlinear fit converged to the same values as the linear fit (tableS1 and S2). Each replicate was first fitted separately and then used to calculate a weighted average (see Tab.S2 for fit parameters).

The measured escape rates versus voltage are plotted in Fig. 5b. For all mutants, we observed slopes that were less negative than for the M2-MspA, indicating the presence of EOF. The slopes for the N90K and N91K mutants were even positive, indicating very strong EOF where *F_EOF_*scales even faster than *F_EP_* with voltage. Adapting Eq. 6 to fit the nonlinear exponent of Eq. 7, we obtained

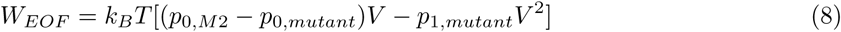

We observe that the *W_EOF_*scales close to linear with voltage for most mutants, with some nonlinearities setting from about 70 mV (Fig. 5c).

Comparing *W_EOF_* at 80 mV we found values of 5.6-12.5 *k_B_T* for the work done by EOF in the various MspA mutants (Fig. 5d). In the following plots, we use the *W_EOF_* calculated at 80 mV unless otherwise specified. For comparison, we also calculated the work done by EOF using a linear fit for both the reference and EOF condition (see Fig. S9, and Tab.S1 and S5) and obtained very similar results.

We find that the strength of the EOF is strongly determined by their location with respect to the nanopore construction. Plotting *W_EOF_*against the radial distance of the charged atoms closest to the nanopore axis for each of the mutants, we found a linear relationship described by the black line in Fig. 5e. This can be understood as the closer a charge sits to the central axis of the nanopore, the more of the nanopore chennel’s volume will be occupied by the Debye layer, resulting in a higher ion selectivity and stronger EOF. The S103K/N108K and the L88K/N108K double mutants showed slightly higher *W_EOF_* than accounted for by the radial distance, which can be attributed to the contribution of the second N108K mutation further away (shown with non-filled markers in Fig. 5e, not included in the fit). This finding is in good agreement with previous work.[47, 6]

We also measured the EOF work for the E_10_N_10_R_x_, x = 7, 8, 9, 10 peptide series in the S103K/N108K nanopore. The EOF work done on the peptide series slightly decreased for lower values of x (Fig. S8). This is a combined effect of the lower charge altering the shape of the trap (Δ*x*) and the shorter peptide length decreasing the viscous shear force the peptide experiences due to EOF.

The EOF work followed the expected pattern of slightly lower energy with decreasing peptide length - see Fig. S8.

We used numerical simulations to compute the energy landscape of the new mutant pores (Fig. S10a). The simulations predicted the order of *W_EOF_*for the various pores in perfect alignment with our experimental results. However, quantitatively, it drastically underestimated the EOF, especially for charge positions further from the constriction (Fig. S10b). While work predicted for the N91K mutant was half of that measured experimentally (5.86 *k_B_T* vs. 12.5 *±* 1.1 *k_B_T*), the predicted work for the L88K/N108K mutant was 21 times lower than the experimental value (0.26 *k_B_T* vs. 5.6 *±* 0.6 *k_B_T*). The S103K mutants profile also followed close to that of the S103K/N108K mutant, indicating that the S103K mutation dominates the energy profile in the simulations (Fig. S10a). The quantitative disparities between the numerical simulations and experimental values likely derive from the fact that the ion distribution is approximated as a continuous field, hence underestimating the thickness of the Debye layer.

While our assay measures the work done by EOF on the peptide, it is also interesting to estimate the force that is exerted on an analyte by the EOF. While we cannot experimentally measure this EOF force directly (since we cannot independently measure the distance to the transition state Δ*x* or the profile of the force), we can quantify the EOF force through a combination of experimental and simulation results (Fig. S11). For this, we used the profile of the EOF force versus position of the peptide along the nanopore’s axis as determined from simulations, see Fig. 5f. Since we realized that quantitatively, the magnitude of the force from the simulation may be incorrect, we used our experimentally measured *W_EOF_* to scale the region corresponding to the *trans*-side exit (Fig. 5f, shaded area). This provided an estimate of the magnitude of the force *F_EOF_* versus position, (Fig. 5f), which appears to be about 5 pN. This is the same order of magnitude as has been estimated on proteins from other experiments in biological nanopores[53, 47], and for the electrophoretic force measured on DNA in solid state nanopores.[31, 50]

Next, we measured the ion selectivity of all nanopores, which is commonly reported as a proxy for the strength of the EOF.[47, 6, 38] We experimentally deduced the ion selectivity of the nanopores by measuring the reversal potential for a 2 M *cis*:500 mM *trans* asymmetric KCl buffer and then calculating the permeability ratio from the Goldman-Hodgkin-Katz (GHK) equation (supplementary section S1.5). Fig. 5g plots our experimental *W_EOF_* values against the measured ion selectivity (see Tab.S6 for values). Note that the point for the M2-MspA lies on the x-axis, as the work done in this mutant is by definition zero (*W_EOF,M_*_2_ = 0; Fig. 5g, blue star). We find a strongly nonlinear dependence of *W_EOF_*versus ion selectivity, suggesting a saturating relationship where *W_EOF_* reaches a limiting value at the highest ion selectivities (the black line denotes such a fit). Given this, the relative change in work and force is thus larger at lower values of ion selectivity than at higher ones.

Using the measured ion selectivities, we also estimated the water flux through a nanopore, following Gu *et al.*[27] who described this as

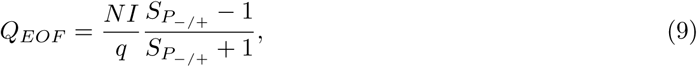

where *Q_EOF_* is the net flux of water molecules through the nanopore per unit of time, *N* is the number of water molecules in the hydration shell of the ions that carry the ion current (commonly estimated to be 6 for the primary hydration shell), *I* is the open state current, *q* the elementary charge, and *S_p_−_/p_*+ is the ion selectivity. Figure 5h plots the experimentally deduced *W_EOF_* versus the estimated *Q_EOF_*obtained from Eq. 9 (see also Tab.S7). Again, we observed a nonlinear relationship. The nonlinear dependence of *W_EOF_*found for both the ion selectivity and *Q_EOF_* could stem from various reasons: (i) The equation from Gu *et al.* may overestimate the water flux from the ion selectivity at higher values. (ii) The charge localization and ion selectivities may be different in the experimental conditions for obtaining *W_EOF_*and the experimental conditions for measuring the reversal potential (e.g. salt concentration and voltage)[54] And (iii) The water flux might not directly translate to the EOF force experienced by the peptide.

We also note that by extrapolating the fit line in Fig. 5g to 1, and Fig. Fig. 5h to 0, it is possible to get a rough estimate of the work done by the EOF in the M2-MspA, otherwise not captured by our method (Fig. S12). For the two graphs we obtain EOF estimates of 1.7 *±* 1.1 *k_B_T* and 1.9 *±* 0.3 *k_B_T*, respectively.

For completeness, we also present the same data with the ion selectivity estimated using the Nernst equation (Fig. S13 & supplementary section S1.5). Overall, we conclude that ion selectivity is not a good proxy for estimating EOF, except for cases of weak EOF.

The fits of the escape rates versus voltage (Fig. 5b) did not only yield the EOF work, but also provided estimates for *k*_0_, i.e., the projected rates at *V* = 0. These *k*_0_ rates differ remarkably among the various mutants. Figure S14 depicts a barplot of the projected rates at *k*_0_. The nanopore with the highest *k*_0_ was M2-MspA, followed by the mutants that line up in order of decreasing distance between the charge and the *trans* side as seen in Fig. 5i. Based on this, we suggest that these shifts can be attributed to the electrostatic repulsion between the charges in the pore and the positively charged end of the peptide. For example, we see that this effect is stronger for the S103K mutant compared to the N91K mutant. As the positive end of the peptide traverses the constriction site in the N91K mutant (where the mutation is situated in the middle of the positive segment), the repulsive force might expel it towards either the *cis* or *trans* side. In the S103K mutant, however, the charge is placed just below the constriction. Consequently, the electrostatic repulsion will push the positive segment back into the constriction site where the electric field and also the EP force are the strongest, further decreasing the escape rate.

To check if our method was applicable in negatively charged MspA variants as well, we also measured the WT MspA (i.e. the actual MspA from Mycobacterium smegmatis, from which M2-MspaA was derived for nanopore sequencing). The WT MspA carries negative charges around the constriction site at the D90, D91, and D93 positions.[25] In addition, it carries three additional charges at the pore rim and in the vestibule, which also were mutated to positive charges in the creation of the M2-MspA (D118R, RD134R, and E139K; Fig. S15a).[12] Now measuring at positive voltage, we found a strong EOF running in the *trans* to *cis* direction, which expelled the peptides back into the *cis* reservoir (Fig. S15b). Accordingly, we chose to instead measure the EOF at negative voltage, where the EOF is still running *cis* to *trans*. Here the peptide is entering the nanopore in the opposite orientation (positive-charged end first). For comparison, the M2-MspA reference was also re-recorded at negative voltages. With decreasing (i.e. more negative) voltage, the WT MspA showed a sharp increase in escape rate, indicating a strong EOF pushing the peptide out of the pore’s *trans* side (Fig. S15c). Calculating *W_EOF_*, we found it to even exceed the EOF value for the N91K mutant, the positive mutant with the highest *W_EOF_* (Fig. S15d). We found *k*_0_ for the WT MspA to be the lowest of any nanopore tested (Fig. S15e). Gratifyingly, the *k*_0_ of the M2-MspA at negative voltage was also found to be similar to that at positive voltage. This also supports the above hypothesis of electrostatic repulsion underlying the pattern in *k*_0_. The results of the WT MspA show that our assay is also applicable and consistent at negative applied voltages.

## 3 Discussion and conclusion

In this work, we developed a method to experimentally measure the work performed by EOF in nanopores, using a dipolar peptide and a reference nanopore with negligible EOF. Our approach provides a quantitative metric of the strength of the EOF, and therefore constitutes an improvement over previous experimental techniques which addressed EOF qualitatively.

We measured six different MspA variants and found the work to be in the range of 5.6-16.7 *k_B_T*. The primary structural predictor of the strength of the EOF in nanopores was found to be the radial distance between the charged residues in the inner pore lumen. Numerical simulations provided insight into the underlying energy landscapes and the forces, and allowed us to estimate the EOF force on the analyte to be approximately 5 pN for the S103K/N108K MspA mutant.

Comparing our experimentally obtained work to the nanopore’s ion selectivity and the water flux estimated from it, we found a strongly nonlinear relationship, indicating that the ion selectivity or water flux does not scale linearly with the EOF force experienced by the peptide. The analysis of *k*_0_ indicated that electrostatic repulsion between the pore and the analyte influences a proteins translocation through the nanopore independently of the EOF.

Our novel method is likely very well generalizable, as already illustrated by the fact that we could apply it to WT MspA which has the opposite charge from the M2 mutants. We expect that it can be applied to a wide variety of other pores beyond MspA, although it may be convenient to redesign the peptide probe for specific applications.

With the quantification of EOF that our new method enables, we can for the first time directly compare the EOF between different conditions. While we here mainly have investigated the effects of different charge positions, we expect that it will be possible to study the effects of salt gradients (see supplementary section S1.4), buffer composition, and geometry with our method. This will help in the future design of nanopore detection systems, especially nanopore protein sequencing systems in which EOF does not just help capture the analyte, but actively keeps it stretched as it ratchets through the pore.

Taken together, our method allows a way to directly compare the work performed by the EOF between different nanopores or conditions, which will help in the rational optimization of EOF for its application in nanopore sensing, sequencing and beyond.

## 4 Acknowledgments

We thank Bauke Albeda, Xiuqi Chen, Justas Ritmejeris, Luning Yu, Verena Rukes, and Jaco van der Torre for discussions, and Eli van der Sluis for in house expression of the M2-MspA and the S103K/N108K MspA mutants. Most M2-MspA mutants used in this study were a kind gift from Giovanni Maglia. We thank Alwin Jansen and Giovanni Maglia for the advise on ion selectivity measurements. This work was funded by the NWO XL grant ProPore and in part supported by Swedish Research Council (Vetenskapsrådet) under Grant 2025-05333, Magnus Bergvalls Stiftelse in Sweden under Grant 2025-233, and ÅAForsk Foundation in Sweden under Grant 26-7. Finally we acknowledge Oxford Nanopore Technologies for funding of this work under a research agreement and for supplying us with the wildtype MspA nanopore.

## 5 Methods

### 5.1 Chemicals

The E_10_N_10_R_10_ and E_10_(SG)_3_SKSG_10_model peptides were obtained from Biomatik. The other peptide variants were obtained from GenScript. The M2-MspA and S103K/N108K MspA were expressed in house as in Wang *et al.*[51]. The other mutants were a kind gift from Giovanni Maglia. The WT MspA was a kind gift from Oxford Nanopore Technologies.

### 5.2 Nanopore measurements

For the nanopore experiments, we used a home-built U-tube device with an aperture fashioned from heat-shrink tubing. AgCl-electrodes were used. Both *cis* and *trans* resorvoirs were filled with symmetric buffer containing 900 mM KCl, 10 mM HEPES at pH 8.0, unless specified otherwise. A bilayer was established using DPhPC lipids from Avanti Polar Lipids (SKU: 850356C), mixed with hexadecane. Using National Instruments X series DAQ operated with custom LabVIEW software, a cross-membrane voltage was established, typically 100 mV for the nanopore insertions. About 0.5 µl of 1 µM of the MspA nanopore variant was added to the *cis* reservoir. A correct nanopore insertion was characterized by a current of approximately 180 pA at 100 mV, although the exact value differed between variants. An IV-curve was taken for each nanopore by measuring the current at every 20 mV between −200 and 200 mV for M2-MspA, or every 10 mV between −100 and 100 mV for charged variants.

### 5.3 Trapping experiments

The model peptide was added to the *cis* reservoir, typically 1 µl of 100 µM, but the amount was adjusted based on the capture rate. Trapping times were recorded in traces taken at every 5 mV with a minimum of 150 events recorded per voltage level. The minimum recordable voltage level was set by an insufficient peptide capture, or, in some cases, by a transition to a *cis*-side dominated escape regime. The maximum recordable voltage level was set by (i) the trapping time becoming too long for efficient recording, (ii) the trapping time becoming too short for correct characterization, (iii) gating becoming to severe at higher voltages, or (iv) trapping of multiple peptides simultaneously (cf. supplementary section S1.1). Most recordings were made at 50 kHz sampling frequency with a 10 kHz Bessel filter. If the trapping time was very short, the recordings were made by sampling at 250 kHz with a 100 kHz Bessel filter.

### 5.4 Ion selectivity measurements

Ion selectivity was recorded as in Baldelli *et al.*[6]. We used 3% agarose 3 M KCl salt bridges between the electrodes and the reservoirs. Both reservoirs were filled with a buffer consisting of 500 mM KCl, 10 mM HEPES, pH 8.0. After nanopore insertion, the voltage offset was corrected to 0 V and an IV-graph was recorded. The *cis* reservoir was then perfused with 2 M KCl, 10 mM HEPES, pH 8.0. The reversal potential was recorded by measuring the current at every 5 mV between −40 and 40 mV. For calculating the ion selectivity, KCl activity values were obtained from Marques *et al.*[36]

### 5.5 Data analysis

The ion current data was typically downsampled to 10 kHz. A polynomial function was fitted to the open state segments to compensate for any baseline drift. Events were identified when the current dropped below a certain threshold value, which typically set to 60% of the open state current. The last datapoint before the threshold was chosen as the start of the event, and the last local minima was chosen as the end of the event.[44] To obtain the current drop of the events (especially short lived events), the current values were fit as in Balijepalli *et al.*[7] Then the data points that differed from this value by typically more than 10% of the open state current were excluded to remove signals due to electrostatic spikes and gating.

For each voltage, the trapping times were binned into a histogram and fitted through a *χ*^2^ least-squares fitting to a single-exponential decay model, to determine the mean and its standard error of the mean.[34, 26] The trapping times were converted to rates, plotted against the voltage, and fit to the expressions described in the main text. Replicates were fitted individually and then used to calculate a weighted average for plotting and calculating the work.

### 5.6 COMSOL simulations

Numerical simulations of the MspA nanopore were implemented on COMSOL Multiphysics 6.1[19] with a two-dimensional axial symmetrical domain. The simulation included the fluid domain, the lipid membrane, the MspA nanopore, and the peptide. The relative permittivity was set to 80, 5[52], and 3.23[3] for water, lipid bilayer, and protein (MspA and peptide), respectively. The profile of the MspA nanopore was a copy from ref [12] with the dimension in scale, and the corresponding positive and negative charge on the nanopore inner wall was set at respective positions. Instead of point-wise charge, we adopted a uniform surface charge band at each charge position with respective charge density. The peptide was approximated by a cylindrical column with a diameter of 0.6 nm, located at the central axis of symmetry of the system. The ion distribution and movement in an electrolyte were governed by the Nernst–Planck equation, the electric potential distribution was described by the Poisson equation, and the water flow was determined by the Navier–Stokes equations. The Transport of Diluted Species module (Nernst–Planck equation), the Electrostatics module (Poisson equation), and the Laminar Flow module (Navier–Stokes equations) were incorporated and fully coupled in the simulation. The electrolyte was KCl with certain concentrations, and the diffusivity of K^+^ and Cl*^−^* were 1.96*×*10*^−^*^9^ and 2.04*×*10*^−^*^9^ m^2^ s*^−^*^1^, respectively.[2]

The respective diffusivity was then determined through the Einstein relation. During the simulation, the peptide was moved downwards manually with a step of 1 nm in the middle part and 2 nm in the start and end parts of the movement. At each step, the steady solutions of corresponding physics fields were found with different bias voltages, in agreement with that used in our experiments. The forces from electrophoresis, *F_EF_*, and electro-osmotic flow, *F_EOF_*, on the peptide were obtained by integrating electromagnetic stress tensor and shear stress tensor on the surface of peptide, respectively. To get the ionic current, the ion flux was integrated on the cross-sectional area located at narrowest restriction of the nanopore.

## Authorship contribution

T.E. and C.D. conceptualized the study. T.E. designed and performed the experiments, developed the analysis code and analyzed the data. C.W. performed the COMSOL simulations. T.E. wrote the manuscript, and C.W. and C.D. reviewed and edited it. C.D. supervised the project.

## Data availability

The data supporting the findings of this study have been deposited in Zenodo under accession code https://doi.org/10.5281/zenodo.24101689. The data will be made publicly available upon publication.

