## Supplementary Information for "Quantifying electro-osmotic flow in mutant MspA nanopores"

November 6, 2026

### S1 Supplementary Notes

#### S1.1 Multiple peptide trappings in the same MspA

When measuring dipolar peptides in the M2-MspA nanopore we discovered that multiple peptides could be trapped in the pore simultaneously.

The first peptide we tried to measure was $E_{10}(SG)_3SKSGR_{10}$, consisting of 10 glutamates, 5 SG repeats with the glycine in the fourth repeat replaced by a lysine, and 10 arginines (Fig.S1a). Upon application of this peptide to the *cis* reservoir filled with 400 mM KCl and an applied potential of 150 mV over the nanopore, we observed a series of three subsequent current drops, which was very reproducible (Fig.S1b). First, the current dropped from the open state ($\sim$110 pA) to $\sim$70 pA, which then continued to fall to $\sim$35 pA and finally to $\sim$10 pA. The peptides did not spontaneously exit the pore, but were expelled by briefly reversing the voltage. We conjectured that the subsequent current drops might be due to multiple peptides being trapped in the pore simultaneously due to the thin, central SG-segment in the $E_{10}(SG)_3SKSGR_{10}$ peptide - see Fig.S1c.

We reasoned that for a peptide with a thicker central segment, such as 10 asparagines ($E_{10}N_{10}R_{10}$; see Fig.S1d), three peptides would not be able to fit in the pore simultaneously. Adding the $E_{10}N_{10}R_{10}$ peptide to the *cis* reservoir under the same conditions as the previous peptide, we indeed observed only two distinct current levels in each trapping event. Because the $E_{10}N_{10}R_{10}$ peptide showed two current levels as opposed to three, we concluded that only two peptides could fit in the nanopore at a time (Fig. S1f).

To make sure the different levels was not just one peptide changing conformation, we measured the $E_{10}(SG)_3SKSGR_{10}$ peptide at 10, 2, and 0.2 nM concentrations in 400 mM KCl at 180 mV (Fig.S1g). At lower concentrations, not only did we see the initial capture time increase, but the time to the second and third current drops increased considerably as well. If there was only one peptide changing conformation, the timing of the subsequent current drops would instead not have been influenced by the concentration. This further confirms that several peptides trap in the pore.

Multiple trappings occurred primarily at higher applied voltages. We measured the $E_{10}N_{10}R_{10}$ peptide across a range of voltages and found that the fraction of events that trapped two peptides was not be explained by capture rate alone. Fig.S1h shows example traces of the peptide trapping at voltages 115, 120 and 125 mV in 900 mM KCl. At 115 mV, most events trap a single peptide, which then exits the pore spontaneously. At 125 mV, however, approximately half of the observed events trap two peptides which had be expelled by reversing the voltage. Investigating the fold change of both the capture rate and the fraction of events trapping two peptides (Fig.S1i) we observed a drastic increase in the fraction of events trapping two peptides, much larger than the increase of the capture rate, starting around 110 mV. This indicates that the increase in events trapping multiple peptides cannot be explained by the capture rate alone. Both the capture rate and the rate for multiple peptide events increased exponentially. As to why the trapping of two peptides occurs more frequently than the capture rate with respect to voltage, we conjecture that this relates to how the first peptide influences the electric field or the charges of the peptides induces EOF, making the capture of the second peptide different.

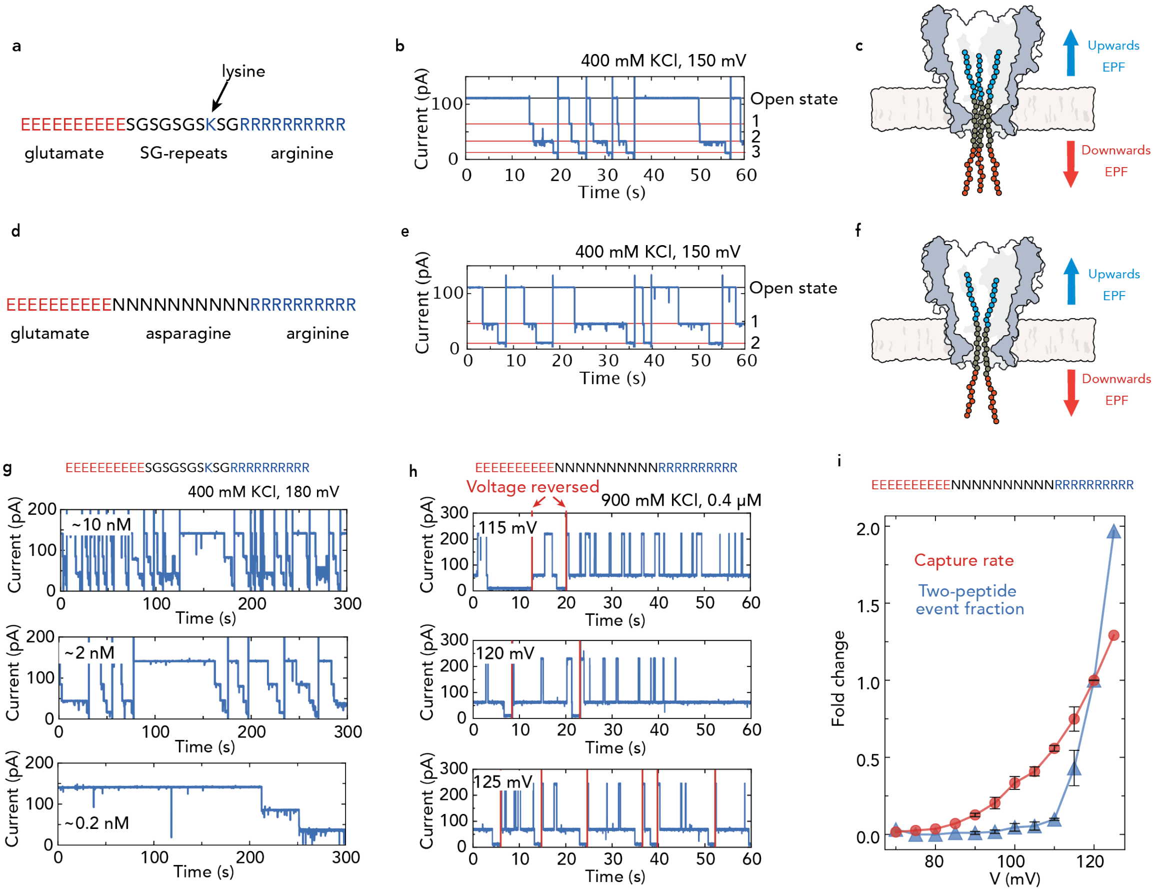


Figure S1: **Description of events where multiple peptides trap into one nanopore.** a) Sequence of the peptide $E_{10}(SG)_3SKSGR_{10}$, containing a sequence of 10 glutamates, 5 SG-repeats with 1 glycine replaced with a lysine, and ten arginines. b) Typical trace of the $E_{10}(SG)_3SKSGR_{10}$ peptide trapping in the nanopore when added to the *cis* reservoir. The peptide did not exit spontaneously, so the voltage was momentarily flipped in between events. In all capture events we see subsequent current drops occurring in steps of three, with levels that are very reproducible between events. c) Cartoon of the system with three $E_{10}(SG)_3SKSGR_{10}$ peptides trapped. We conjecture that due to the narrow SG-repeat segment, up to three peptides can be trapped in the pore simultaneously. d) Sequence of the second peptide $E_{10}N_{10}R_{10}$, containing a series of 10 glutamates, 10 asparagines and 10 arginines. e) Typical trace of the $E_{10}N_{10}R_{10}$ peptide trapping in the nanopore when added to the *cis* reservoir. The peptide did not exit spontaneously, so the voltage was momentarily flipped in between events. Unlike in (b), we see only see two subsequent drops in each event. f) Cartoon of the system with $E_{10}N_{10}R_{10}$. Because of the bulkier asparagine segment in the middle, only two peptides fit in the pore simultaneously. g) Traces of capture events for the $E_{10}(SG)_3SKSGR_{10}$ peptide at ∼10, ∼2, and ∼0.2 nM concentrations. Both the time between events and the time to the second current drop within an event became longer with decreasing concentration, suggesting that multiple peptides are trapped in the pore. Voltage was reversed after the third subsequent current drop to expel the peptides from the pore. h) Example current traces for the $E_{10}N_{10}R_{10}$ peptide at 115, 120 and 125 mV in 900 mM KCl. At higher voltages, double trappings make up a larger fraction of the total events. i) Fold change for the capture rate (red) and the fraction of events trapping two peptides (blue) for the $E_{10}N_{10}R_{10}$ peptide. We see a very sharp rise in the fold change of the fraction of events trapping two peptides at 110 mV, surpassing that of the capture rate. The fold change is calculated from up to three replicates and error bars indicate standard error of the mean . Markers without error bars were only measured in one replicate.

#### S1.2 Estimation of entropy for simulations

We estimated the entropic contribution to translocation of the 30 amino acid peptides through MspA as a coarse-grained conformational free-energy change.[7] The maximum entropic energy change of stretching a peptide can be expressed as follows, for a Gaussian-chain:

$$\Delta F_{\text{max}} = \frac{3k_{\text{B}}TR^2}{2Nb^2}, \tag{S1}$$

where $R$ is the assumed end-to-end distance of the confined peptide, $N$ is its number of Kuhn segments, and $b$ is the Kuhn length [6]. To construct the position-dependent profile, we assumed a constant contribution per unit confined length in the pore, such that

$$\Delta F(z) = \Delta F_{\text{max}} \frac{\ell_{\text{conf}}(z)}{L_{\text{c}}}, \tag{S2}$$

where $l_{conf}(z)$ is the length of peptide in the pore and $L_c$ is the total peptide contour length. Accordingly, the free energy increases during insertion, reaches a short plateau when the entire peptide enters, and decreases as the peptide exits and recovers conformational freedom (Fig.S5)b.

#### S1.3 Nonlinear scaling of EOF in the S103K/N108K and L88K/N108K nanopores

In order to investigate the nonlinear scaling effect explained in the main text, we measured the peptide series $E_{10}N_{10}R_x$, x = 7, 8, 9, 10, in the S103K/N108K nanopore mutant at 900 mM KCl. The rates were fit to $\ln(k) = \ln(k_0) - p_0V - p_1V^2$. Fig.S2a shows a plot of the rate versus voltage. For lower $x$ values, we see the escape rate increase, as the *cis*-directed EP force decreases. Due to the lower EP force, the slopes initially also become more positive for lower values of x. Accordingly, the maximum rate is shifted towards higher voltages (red tacks in Fig.S2a). The curvature of the fit is given by $p_1$ and can be seen in Table S2. $p_1$ is comparable between the different peptides, but might decrease slightly for lower x-values.

If the nonlinear effect was related to lower ion concentration localization with increasing voltage (as discussed in the main text), we reasoned that the nonlinear behavior in the L88K/N108K mutant would be different than in the S103K/N108K mutant due to the different charge position. Therefore, we measured the $E_{10}N_{10}R_8$ peptide in the L88K/N108K mutant at 900 mM KCl as well. A plot of the rate versus voltage can be seen in Fig.S2b, where the brown line denotes the L88K/N108K mutant and the blue line is the same peptide in the S103K/N108K mutant for comparison. The nonlinear effect was less pronounced in the L88K/N108K mutant (Fig.S2b and $p_1$ in Tab.S2). If the hypothesis of the decreased ion concentration localization is correct, we can explain the the weaker nonlinearity by the charge placement in the L88K/N108K mutant sitting further from the pores central axis. Accordingly any change in the Debye layers thickness would have a smaller effect.

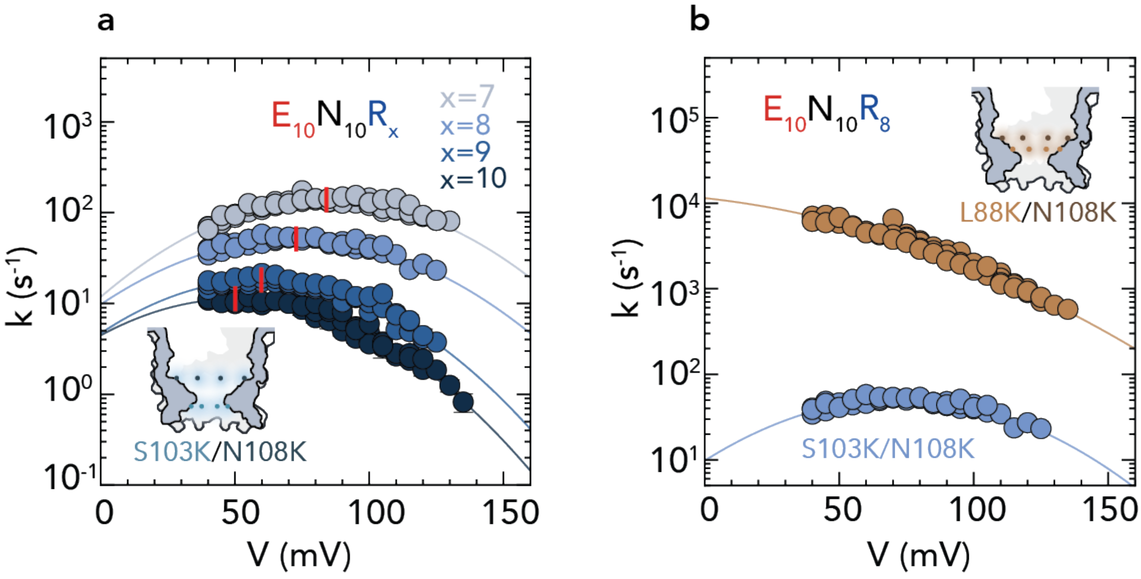


Figure S2: **EOF work on $E_{10}N_{10}R_x$, x = 7, 8, 9, 10 peptide series in the S103K/N108K and L88K/N108K mutant nanopores.** a) Rate vs. voltage for peptides x = 7, 8, 9, 10 in the S103K/N108K mutant. Lines describe $\ln(k) = \ln(k_0) - p_0 V - p_1 V^2$. Red marks note the maximum rate for each fit. b) Rate vs. voltage for the $E_{10}N_{10}R_x$, x=8 peptide in the L88K/N108K mutant (brown) and the S103K/N108K mutant (blue). In addition to the lower EOF in the L88K/N108K mutant (more negative slope), we see it exhibit less of the nonlinear effect observed in the S103K/N108K mutant. Lines describe $\ln(k) = \ln(k_0) - p_0 V - p_1 V^2$. All pores are measured at 900 mM KCl. Error bars indicate standard error of the mean where visible.

#### S1.4 Effects of salt concentration on $W_{EOF}$

The salt concentration has two competing effects on the EOF. At very low salt concentrations, the Debye layer is very thick, and consequently the Debye layers from charges at different positions around the nanopore channel will overlap. As a result, the counterions in the Debye layers are not sufficient to fully screen the surface charge, so the total amount of counter ions is smaller and the EOF will be weaker. When the salt concentration is increased to the point where the Debye layers no longer overlap, the velocity of the EOF will reach its maximum value. Increasing the salt concentration further will have no effect on the EOF's velocity, but will alter the volume of the driving force of EOF as the Debye layer continues to thin. See Section 9.3 of Ref. [3].

In short nanochannels such as nanopores, a backward flow or vortex can be created due to pressure buildup at the mouths of the pore as a direct consequence of the EOF. Assume an EOF is running *cis* to *trans* through a nanopore. The EOF accelerates the flow above the nanopore and retards it below the nanopore. Consequently, due to the incompressible nature of water a pressure gradient will build up across the nanopore. As the EOF increases with voltage, so will the strength of this pressure gradient. At high salt concentrations, more ions move through the pore in the EOF. As the Debye layer continues to thin, the pressure gradient might create a considerable reverse flow through the bulk region of the nanopore, decreasing the net water flow. [5, 8] Together, these two effects suggest a maximum EOF at the salt concentration where the Debye layers are fully developed but before the obvious retardation from the buildup pressure sets in.

To investigate this relationship, we measured the EOF work on the $E_{10}N_{10}R_8$ peptide at 500, 600, and 900 mM, and 2 M KCl. Because the Debye layer is already fully developed at 500 mM KCl, our concentration series only captured the decrease in the EOF due to the pressure gradient. Fig.S3a shows the escape rate versus the voltage at the different salt concentrations for the M2-MspA, to be used as references in calculating the work. The rates of the 500, 600, and 900 mM salt concentrations are remarkably similar, but the rates for the 2 M condition are noticeably higher and seem to have a slightly nonlinear dependence on voltage. Therefore, the 2 M concentration was fit with the nonlinear expression unlike the lower concentrations fit with the linear expression. We suggest that the different slope might depend on higher screening that reduces the EP force and keeps the peptide trapped in the pore. Fit parameters for both the M2-MspA nanopore and the S103K/N108K mutant are given in Tab.S3.

In the S103K/N108K mutant, the slopes and the curvature of the 500, 600, and 900 mM salt concentrations are similar to each other, as they were in the M2-MspA. However, unlike in the M2-MspA, their $k_0$ differs by more than one degree of magnitude, with higher values for higher salt concentration. This effect is related to the electrostatic repulsion between the charges in the pore and the peptide's positive end, discussed more extensively in the main text. As the salt concentration increases, so does the screening, decreasing the electrostatic repulsion in traversing the nanopore. Also, unlike the M2-MspA, the rates for the 2 M condition have a more negative slope than the lower concentrations, indicating that the EOF force decreases more rapidly than the EP force for the salt concentrations investigated.

We calculated the EOF work through $W_{EOF} = k_BT[(p_{0,M2} - p_{0,mutant})V - p_{1,mutant}V^2]$, or through $W_{EOF} = k_BT[(p_{0,M2} - p_{0,mutant})V + (p_{1,M2} - p_{1,mutant})V^2]$ for the 2 M concentration. The work done by EOF across voltages (Fig.S3c) and at 80 mV (Fig.S3d) was similar for the 500, 600, and 900 mM KCl concentrations ($\sim 9.5\ k_BT$), but then decreased to about a fourth between the 900 mM and 2 M concentrations ($\sim 2.5\ k_BT$). We therefore conclude that a considerable reverse flow emerges somewhere above 900 mM KCl. These results are similar to those reported by Willems *et al.* where they applied a continuum model and found the maximum EOF in the ClyA nanopore to be at 500 mM NaCl, decreasing slightly until 1 M, and then followed by a sharper decrease above 1 M.[9] Complete values for work are given in Tab.S5.

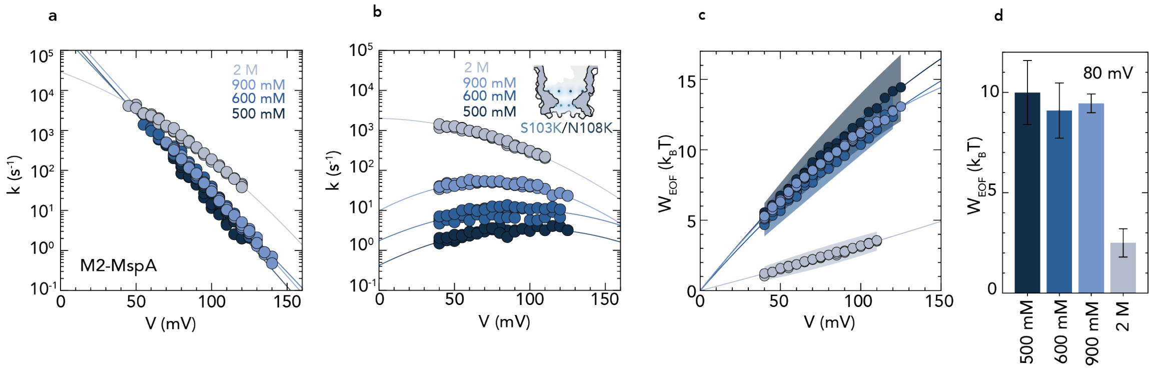


Figure S3: **Effect of the KCl concentration on the EOF work in the S103K/N108K mutant.** a) Escape rate vs. voltage for the $E_{10}N_{10}R_8$ peptide at different salt concentrations in the M2-MspA nanopore. The lines describe the relationship $\ln(k) = \ln(k_0) - p_0V$ except for the 2 M concentration which is described by $\ln(k) = \ln(k_0) - p_0V - p_1V^2$. Error bars indicate standard error of the mean where visible. b) Same as in (a) but for the S103K/N108K mutant. The lines describe the relationship $\ln(k) = \ln(k_0) - p_0V - p_1V^2$. Error bars indicate standard error of the mean where visible. c) Work done by EOF vs. voltage. Calculated through $W_{EOF} = k_BT[(p_{0,M2} - p_{0,mutant})V - p_{1,mutant}V^2]$, or through $W_{EOF} = k_BT[(p_{0,M2} - p_{0,mutant})V + (p_{1,M2} - p_{1,mutant})V^2]$ for the 2 M concentration. The shaded area indicates standard deviation. d) Work done by EOF at 80 mV. The work is similar at the 500 - 900 mM concentrations, but then decreases drastically at the 2 M concentration. Error bars indicate standard deviation.

#### S1.5 Determination of the ion selectivity

As covered in Zhang *et al.*[10], there are two common equations for estimating the ion selectivity from the reversal potential of nanopores: the Goldman-Hodgkin-Katz (GHK) equation and the Nernst equation. However, these equations make different assumptions about the system and are not trivially interchangeable. The GHK equation assumes a constant electric field, which is not necessarily valid for nanopores, where there is a sharp potential drop at the nanopore's constriction site. The Nernst equation, on the other hand, assumes electroneutrality. Both equations assume constant permeability, which does not hold in practice for highly charged channels and different salt concentrations on the two sides of the membrane.[10]

The GHK equation is given for monovalent ions by

$$V_{rev} = \phi_w - \phi_r = \frac{RT}{F} ln(\frac{P_+ a_r + P_- a_w}{P_+ a_w + P_- a_r}) \tag{S3}$$

where $V_{rev}$ is the reversal potential, $\phi_w$ and $\phi_r$ are the working and reference potentials, respectively, $a_w$ and $a_r$ are the activity on the working and reference sides, respectively, and $P$ is the permeability.

The Nernst equation is given by

$$V_{rev} = \phi_w - \phi_r = \frac{RT}{F}(1 - 2t_+) ln(\frac{a_w}{a_r}) \tag{S4}$$

where $t_+$ is the cation transference number, and $t_- = 1 - t_+$.

In general, the ratio between the transference numbers from the Nernst equation, $t_+/t_-$, and the ratio of the ion permeabilities obtained by the GHK equation, $P_+/P_-$, are not equal. Therefore, we adhere to the best practices proposed by Zhang *et al.* and report the measured values of the reversal potential, the concentrations used for the gradient, and the ion selectivities calculated with both the GHK and the Nernst equations for easier comparison with other work. Ultimately, however, both the GHK and the Nernst equations only give estimates of the selectivity in the nanopores.

#### S1.6 Experimental details for measuring EOF with nanopore tweezers

Below is a list of notes on our experimental procedures to make our method more accessible.

##### S1.6.1 Peptide design

In our current nanopore (sequencing) experiments, we generally use voltages between 80-120 mV when measuring the EOF mutant nanopores, and our peptide probe was therefore designed with this voltage range in mind. $E_{10}N_{10}R_{10}$ was the probe chosen for most of our experiments, but it is not necessarily always the optimal peptide. For example, the trapping times of the $E_{10}N_{10}R_8$ were shorter in all of the nanopores and could have been measured more efficiently. However, due to the wide range of rates, the $E_{10}N_{10}R_8$ might have been difficult or impossible to record in some of the pores with very high escape rates.

Consistent with nanopore tweezer theory, we found that peptides with higher charge were trapped for longer and at lower voltage than less-charged peptides. In our initial exploration of peptides, we also tested the peptides $E_6(GS)_5R_6$, $E_7(GS)_5R_7$, and $E_8(GS)_5R_8$ among others, but found that they trapped at inconveniently high voltages.

While in this proof-of-concept work we have mainly concerned ourselves with a symmetric peptide, an asymmetric dipolar peptide might be preferred for more efficient measurements. For example, by virtue of the local electric field at the constriction site, one would imagine that, for example, an $E_{15}N_{10}R_9$ peptide might increase the capture rate while still having appropriately long trapping times. Another possible improvement could be to make the neutral segment of the peptide bulkier to prevent multiple peptides from simultaneously trapping in the pore at higher voltages.

Of course, the peptide in this work is designed for MspA, and other nanopores will likely pose other design considerations.

##### S1.6.2 Concentrations

We found that the escape of a peptide from the nanopore was not influenced by the bulk peptide concentration, something previously reported by Asandei *et al.*[1] Therefore, the peptide concentration in the *cis* well (containing approximately 50 µl) was adjusted during experiments to adapt to a lower capture rate at low voltages or to minimize multiple trappings at higher voltages. In general, the final peptide concentration in the *cis* well was between 0.01-40 µM. For the EOF-generating mutants, a concentration of 4 µM was usually sufficient due to the EOF-enhanced capture rate.

###### S1.6.3 Recording

The voltage range was measured in 5 mV steps, with either *cis*-side exit or capture rates setting the lower limit, and multiple trappings, inconveniently long trapping times, or pore instability setting the upper limit. For accurate averages of the trapping times, we found that a minimum of 150 events should be recorded at each voltage.

###### S1.6.4 Dealing with multiple trappings

The multiple trappings described in the supplementary section S1.1, occurred at higher voltages (>100 mV for our chosen peptide) and usually required a reversal of the voltage to expel it from the pore. The events containing multiple trappings were considered unusable as they altered the trapping time of the peptides in the nanopore. However, the longer a peptide sits in the nanopore the higher the chance for a second peptide to trap. Therefore, they could not simply be discarded as that would skew the distribution of trapping times.

We found the best way to reduce multiple trappings was to reduce the analyte concentration. However, as noted in section S1.1, the capture time of the second peptide scales differently than the first, so the maximum measurable voltage can only be adjusted by 10-15 mV by this lever.

For the $E_{10}N_{10}R_x$, x = 7, 8, 9, 10 peptide series, we found that decreasing the positive charge and upwards EP force also enabled measurement of the peptides at higher voltages.

###### S1.6.5 Filtering conditions

During data processing a number of filters were applied to assure that we only included good quality events. E.g. to discard events with multiple trapped peptides or to reduce events due to the pore gating. Filter conditions applied during data processing were primarily used to discard events with multiple trapped peptides, and secondarily to reduce events from gating. The filtering process was carried out in four steps: (i) During event recognition, if an event contained time points with negative voltages, i.e. the voltage was flipped to kick out multiple peptides trapped in the nanopore, it was recorded as a bad event. (ii) For each event, the current values were binned into a histograms and peaks were recognized using a gaussian mixture model. If there were more than one current level in the event, it was recorded as a bad event. (iii) Taking the remaining events after the previous two steps, only events within ±0.1 $I/I_0$ of the average event current were used. This was to minimize events from gating or electrical spikes. The removed events were considered bad events. (iv) If the bad events recognized in the three previous steps totaled more that 10% of the total found events, the whole voltage level was discarded from that replicate.

###### S1.6.6 Other observations

During measurements, we observed the capture rate increase considerably after a few minutes. We conjecture this is because the charged peptide interacts with the charged head groups of the lipid used for the membrane (DPhPC), effectively reducing the diffuse environment from 3D to 2D. This is aking to how we use a cholesteral anchor to increase capture rate in our peptide sequencing assay.[2] Another explanation could be that as the peptides translocate, the concentration increases locally on the *trans* side, enabling capture from both sides.

### S2 Supplementary Figures

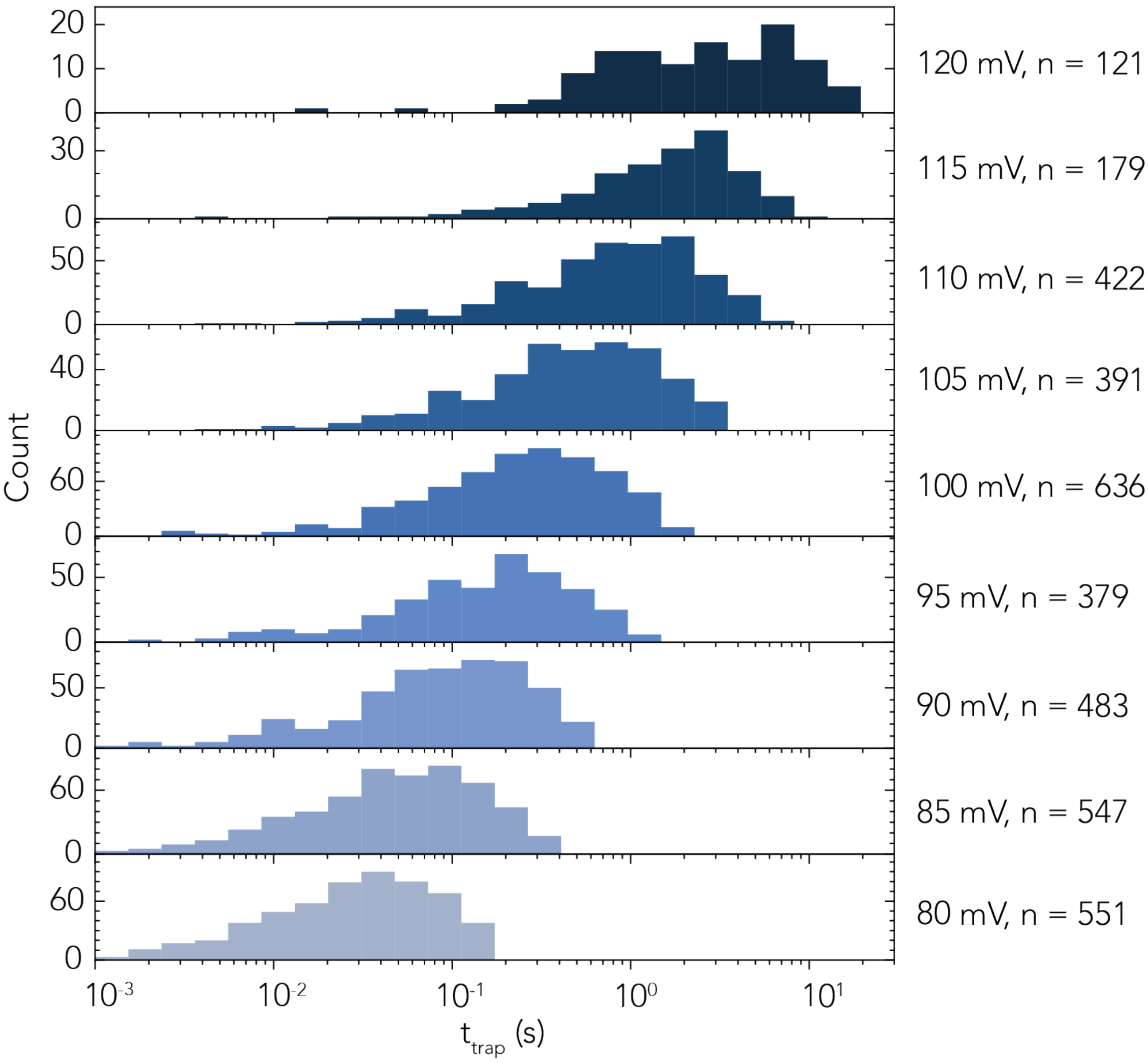


Figure S4: **Histograms showing the trapping times for the $E_{10}N_{10}R_{10}$ peptide for various voltages in M2-MspA.** As the voltage increases, the average trapping time becomes longer. These data are for one replicate.

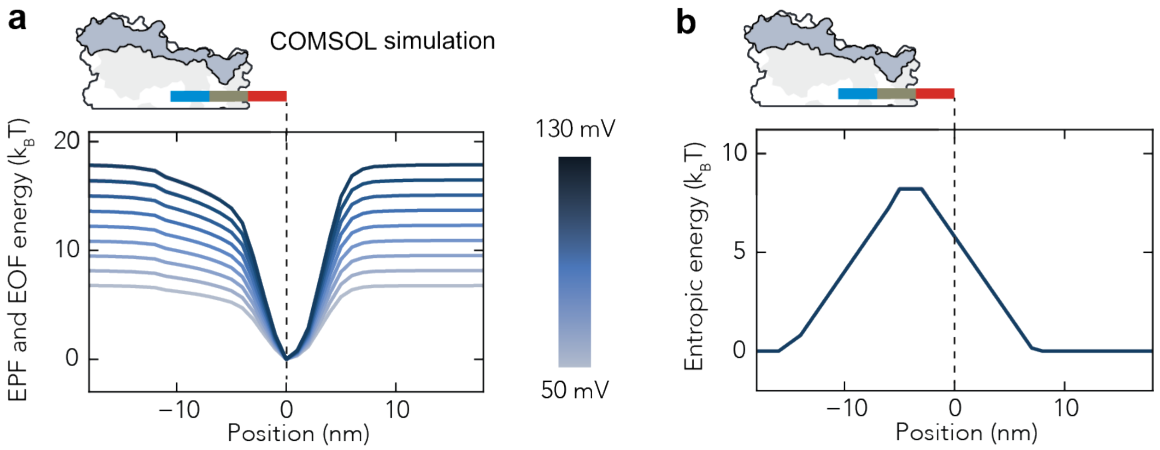


Figure S5: **Contribution of entropy to the energy landscape for the $E_{10}N_{10}R_{10}$ peptide in the M2-MspA nanopore.** a) Numerical simulation of the energy landscape due the EP and EOF on a dipolar rod moved in 1 nm steps along the nanopore axis. b) Entropic energy estimated for the peptide. Information about the simulations and calculation of the entropy can be found in the methods section and supplementary section S1.2, respectively.

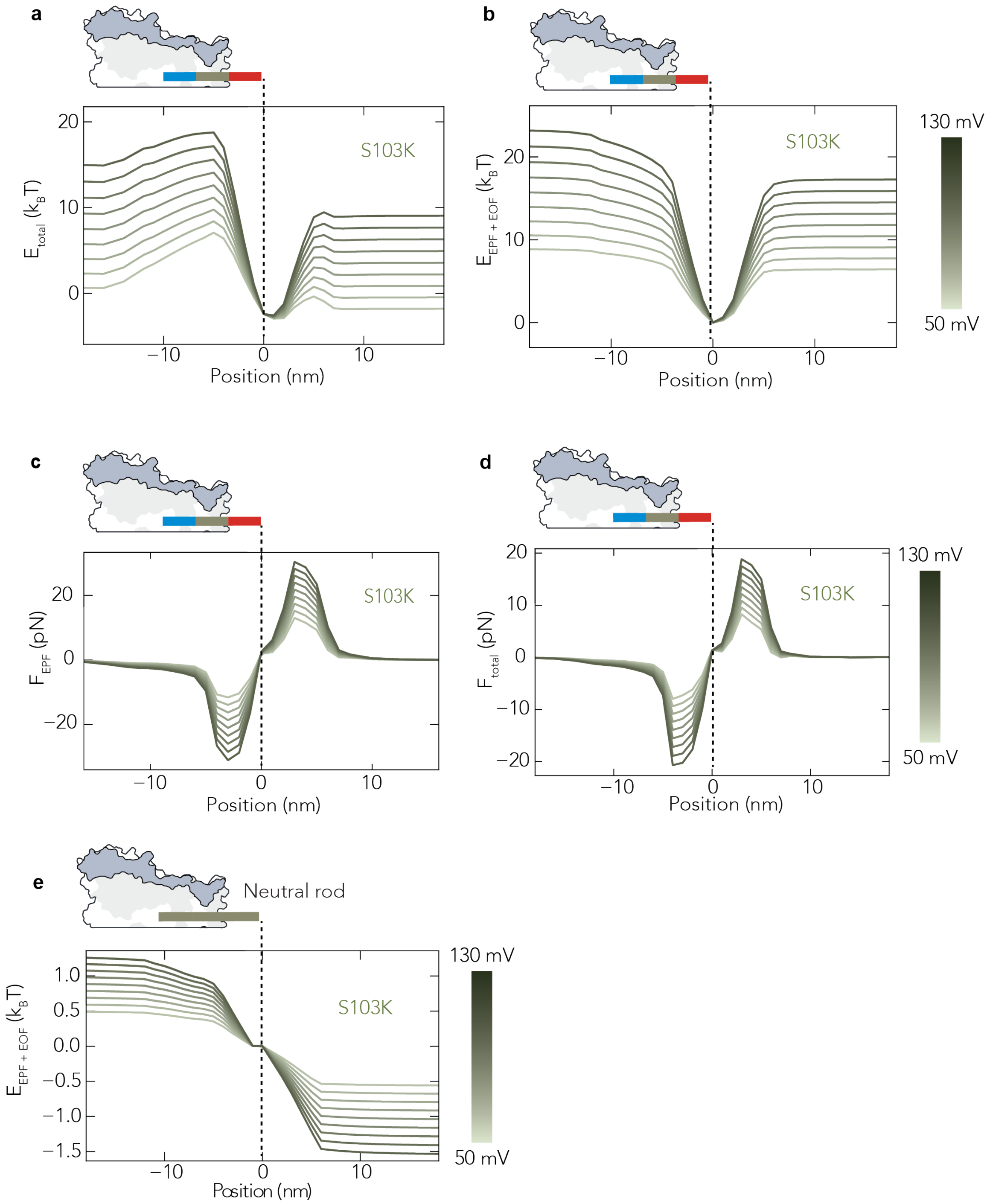


Figure S6: **Numerical simulations of the energy and force for the $E_{10}N_{10}R_{10}$ peptide in the S103K mutant.** a) Energy landscape of the total energy including entropy. b) Energy landscape based on EP and EOF, excluding entropy. c) Electrophoretic force on the peptide. d) Total force $(F_{EP} + F_{EOF})$ on the peptide. e) Energy landscape of a neutral rod in the nanopore, which results only from the EOF generated by the pore. For details about the simulations, see the methods section.

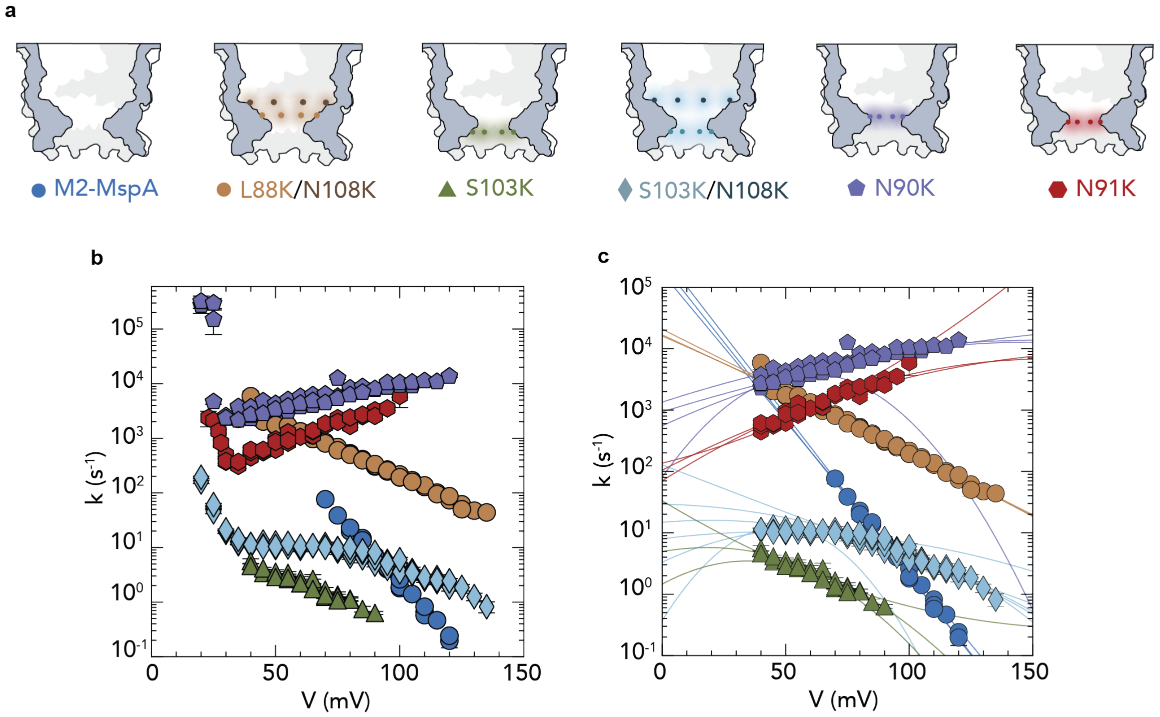


Figure S7: **Data for the $E_{10}N_{10}R_{10}$ peptide trapped in all MspA mutants.** a) Location of the charges in the the various MspA mutants. All mutations are in the M2-MspA background. b) Full set of data points recorded for the mutants. Below 40 mV, the peptides exit towards the *cis* side in a regime dominated by the electrophoretic repulsion between the charged mutations and the positive charges in the peptide for pores S103K/N108K, L88K/N108K, N90K, and N91K. Above 40 mV, the pores exit out of the *trans* side in a regime dominated by EP and EOF forces. Error bars indicate standard error of the mean. c) Individual fits of the replicates. The lines describe the fit $ln(k) = ln(k0) - p_0V - p_1V^2$, with the exception of the M2-MspA which lines describe $ln(k) = ln(k0) - p_0V$. Error bars indicate standard error of the mean.

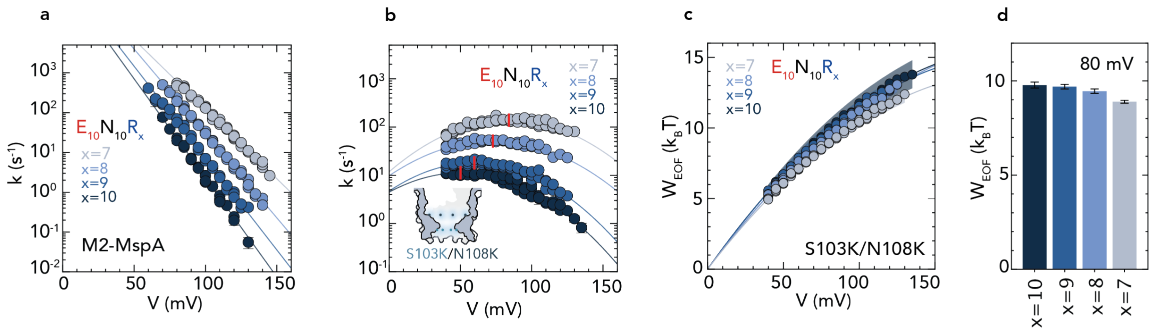


Figure S8: **EOF work for the $E_{10}N_{10}R_x$, x = 7, 8, 9, 10 peptide series in the S103K/N108K mutant.** a) Rates vs. voltage for the peptide series in the M2-MspA nanopore, to be used as references for the S103K/N108K mutant. With decreasing positive charge, the slopes become less negative as a smaller EP force keeps the peptide trapped in the pore. Fits describe the relationship $ln(k) = ln(k_0) - p_0V$. Error bars indicate standard error of the mean. b) Rates vs. voltage for the peptide series in the S103K/N108K mutant. A nonlinear effect is observed in all nanopores and the overall escape rate increases for lower x-values. Fits describe the relationship $ln(k) = ln(k0) - p_0V - p_1V^2$ where the red tack denotes the maximum. Error bars indicate standard error of the mean. c) EOF work vs. voltage obtained through $W_{EOF} = k_BT[(p_{0,M2} - p_{0,mutant})V - p_{1,mutant}V^2]$ for the peptide series. Shaded area represents the standard deviation. d) Work done at 80 mV for the peptide series calculated from the equation in (c). Overall, the work done is very similar between the peptides with a slight decrease for lower x-values. This is in agreement with our model as (i) a shorter peptide will experience less shear force from the EOF, and (ii) the peptides length and charge determines $\Delta x$, the distance between the energy minima and the energy barrier for escape. All measurements were performed in 900 mM KCl. Error bars indicate standard deviation. Fitting parameters are given in Tab.S4 and values for work are given in Tab.S5.

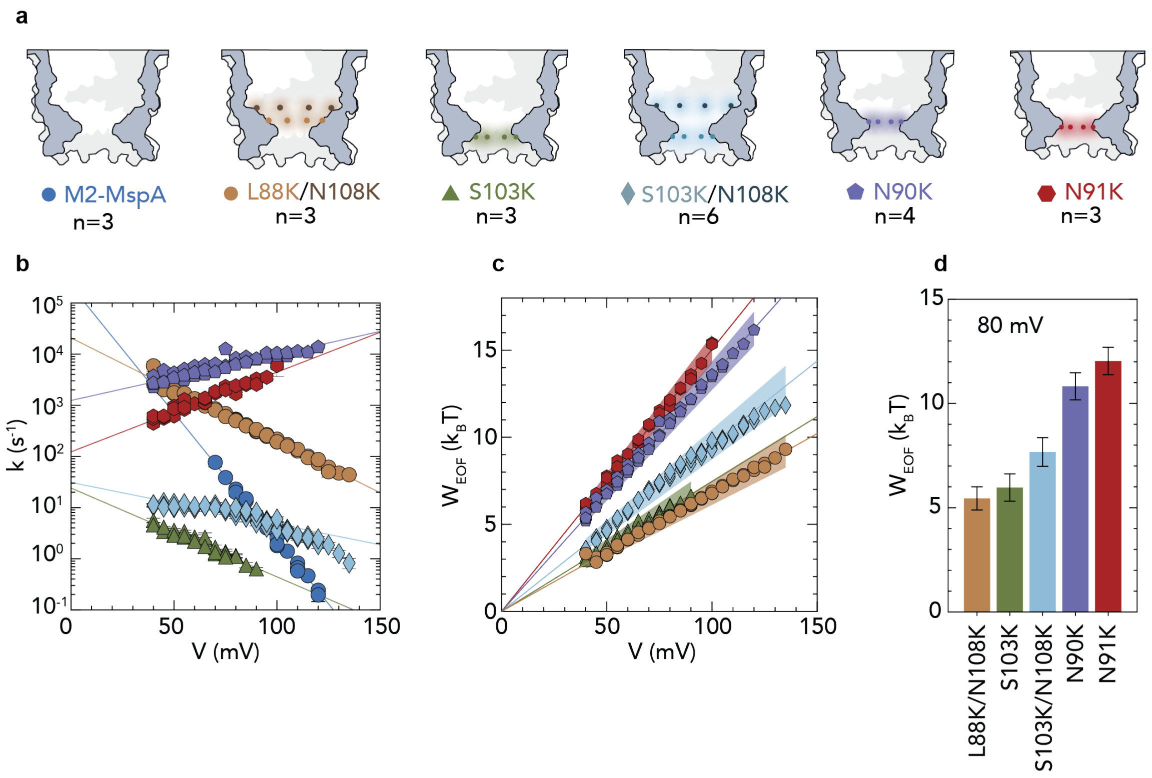


Figure S9: **EOF work for the $E_{10}N_{10}R_{10}$ peptide in the mutant nanopores using the linear** $ln(k) = ln(k0) - p_0 V$ **for fitting both the reference and EOF conditions.** a) Cartoon of the charged position in the different mutants. b) Escape rate vs. voltages for the different mutants. The only mutant noticeably diverging from the linear fit is the S103K/N108K mutant. Error bars indicate standard error of the mean. c) Work vs. voltage for the different mutants calculated through $W_{EOF} = k_B T[(p_{0,M2} - p_{0,mutant})V$. Shaded area indicates standard deviation. d) Work at 80 mV calculated from the same equation as in (c). The work obtained is similar to the curved fit used in the main text for most mutants. Error bars indicate standard deviation. Fitting parameters are given in Tab.S1 and values for work are given in Tab.S5.

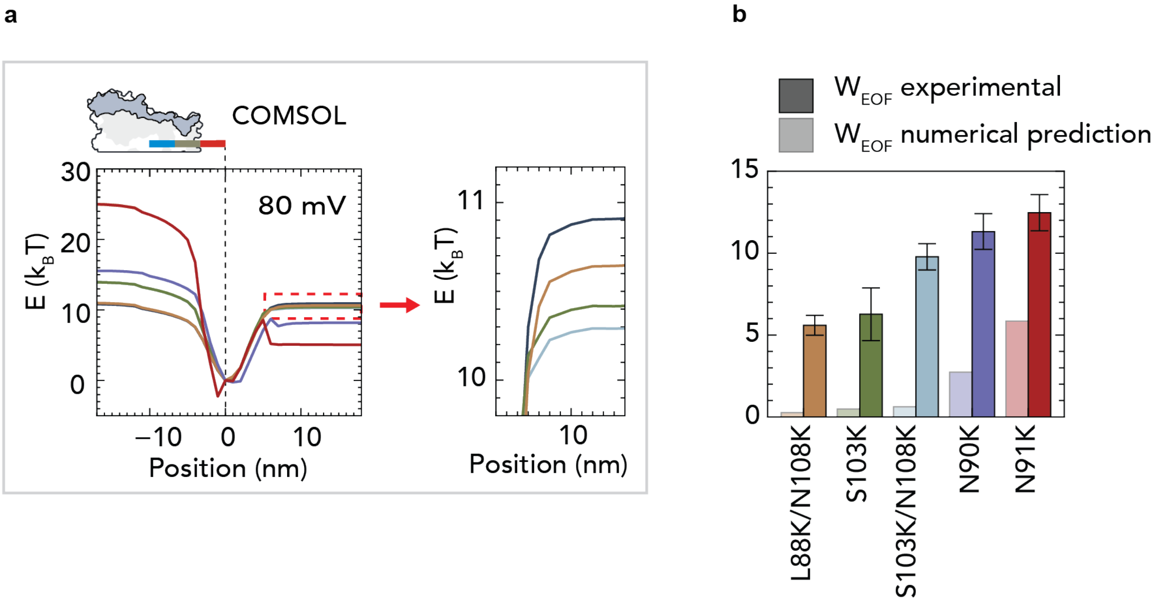


Figure S10: **Numerical simulations of the energy landscape and EOF work in the different MspA mutants.** a) Energy landscape of $F_{EP}$ and $F_{EOF}$, excluding entropy for the various pores. b) Estimated work done by EOF from the numerical simulations (the difference in energy between the mutant pores and the M2-MspA at the 18 nm position in panel (a)), as well as the experimentally measured EOF work. Experimental values are shown in dark colors; numerical estimates in dimmed colors. The numerical simulations appear to systematically underestimate the work done by EOF, with the deviation becoming more severe the further the charges are place from the constriction site. Error bars indicate standard deviation.

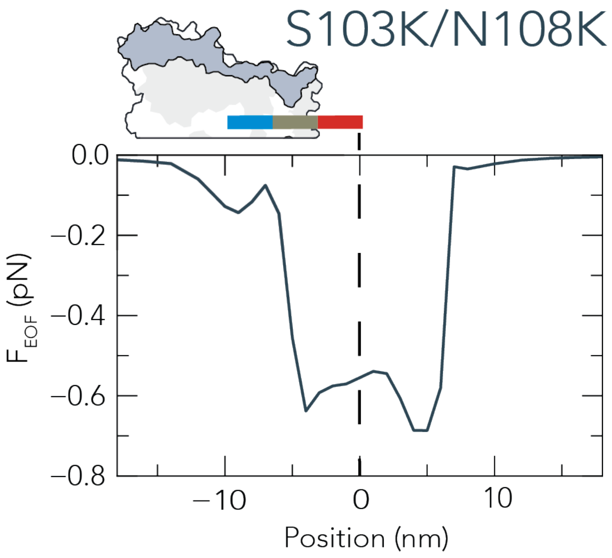


Figure S11: **EOF force on a dipolar rod along the central axis in the S103K/N108K nanopore estimated by numerical simulations.** The absolute force (about 0.6 pN) is an underestimation. The peak at around 5 nm is an artifact from the EOF pushing on the peptides blunt end when it is located in the constriction site.

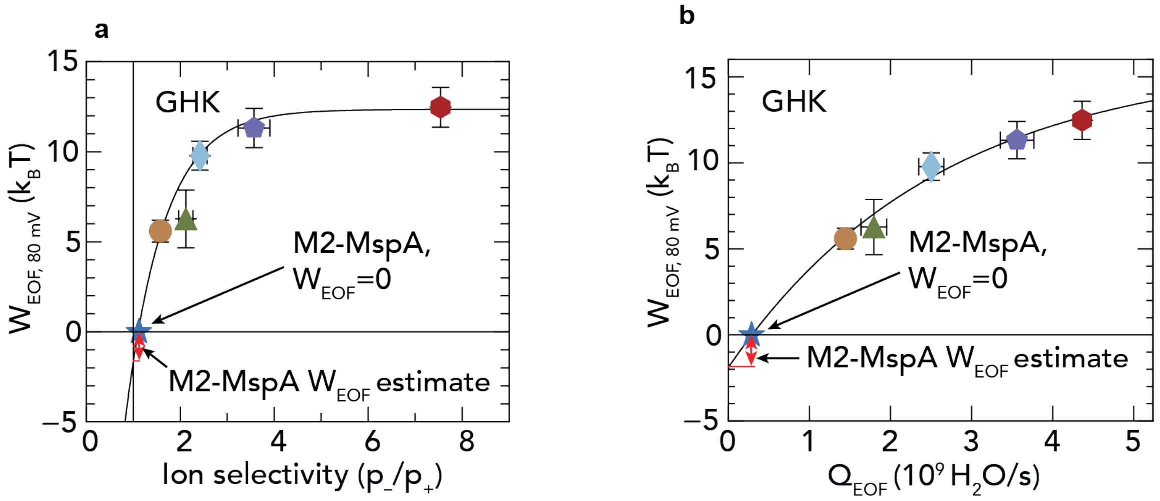


Figure S12: **EOF work estimated in the M2-MspA nanopore.** By extrapolating the fit line in the graph for $W_{EOF}$ vs. ion selectivity in (a) to 1, or the fit for $W_{EOF}$ vs. $Q_{EOF}$ in (b) to 0, we can obtain an estimate of the $W_{EOF}$ in M2-MspA nanopore, otherwise used as a reference. We obtain values of $1.7 \pm 1.1\, k_BT$ and $1.9 \pm 0.3\, k_BT$ for (a) and (b), respectively, which is about a third of the $W_{EOF}$ in the L88K/N108K nanopore (brown) which is the MspA mutant with the lowest EOF. Error bars indicate standard deviation. The line in (a) is fitted to the saturating expression $W_{EOF} = a * e^{-b*S_{p_-/p_+}} + c$, where $a$, $b$, and $c$ are fit parameters, and $S_{p_-/p_+}$ is the ion selectivity. The line in (b) is fitted with the same expression but uses the water flux $Q_{EOF}$ instead of the ion selectivity.

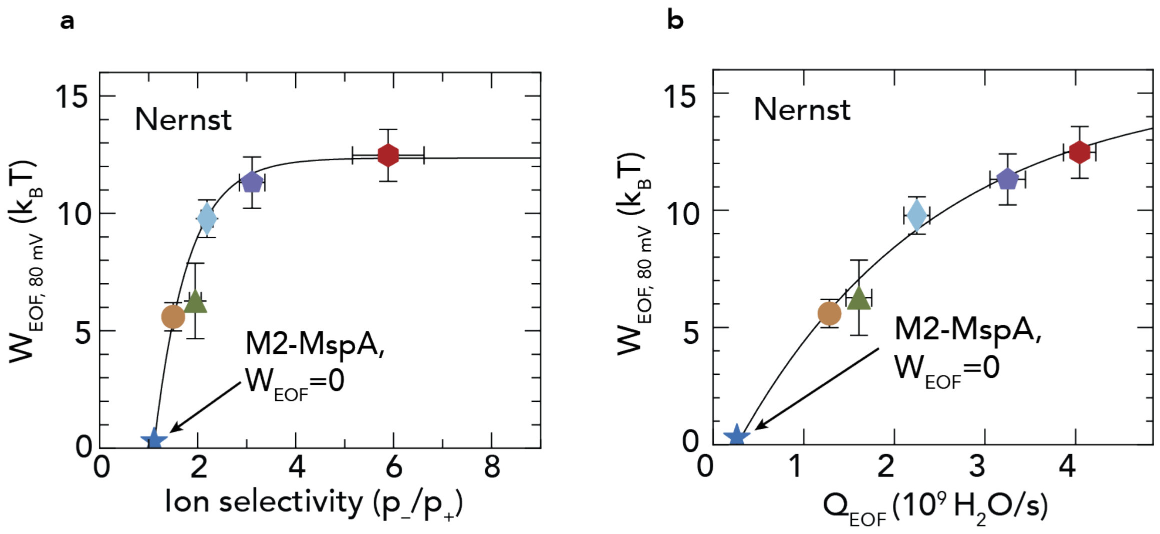


Figure S13: **EOF work versus the ion selectivity and the water flux estimated by the Nernst equation.** a) $W_{EOF}$ versus ion selectivity. The ion selectivity of the different pores was estimated by measuring the reversal potential in the nanopores over a 2 M KCl *cis*: 500 mM KCl *trans* salt gradient, and then using the Nernst equation to calculate the ratio of the ion permeabilities (section S1.5). As the work done by the M2-MspA by definition is zero, it sits on the x-axis at the point of its ion selectivity (blue star). We observe a strongly nonlinear, saturating trend between $W_{EOF}$ and the ion selectivity. The line is fitted to the saturating expression $W_{EOF} = a * e^{-b * S_{p-/p+}} + c$, where $a$, $b$, and $c$ are fit parameters, and $S_{p-/p+}$ is the ion selectivity. b) Work done by the EOF vs. the water flux through the nanopores estimated from the ion selectivity by the equation from Gu *et al.*[4]. Again, we observe a nonlinear trend between the $W_{EOF}$ and the water flux. Like in (a), the M2-MspA sits on the x-axis (blue star). Error bars indicate standard deviation. The line is fitted with the same expression as in (a) but uses the water flux $Q_{EOF}$ instead of the ion selectivity.

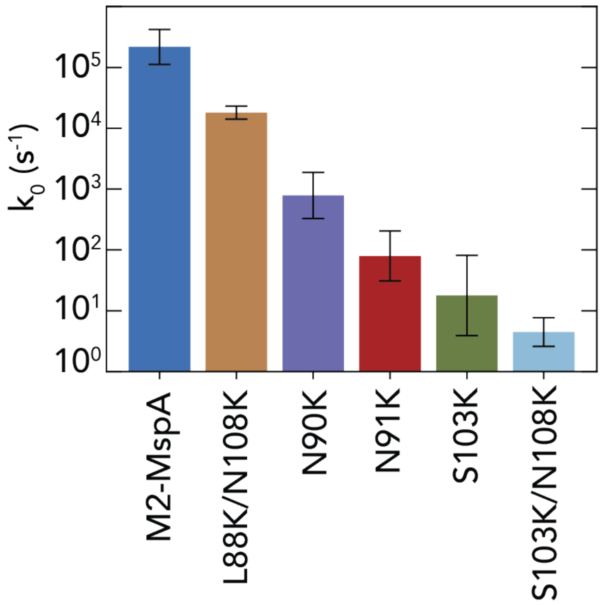


Figure S14: **Intrinsic escape rate $k_0$, i.e., escape rate at $V = 0$, for the different nanopores.** $k_0$ is estimated through a linear fit $ln(k) = ln(k0) - p_0V$ for M2-MspA, and a nonlinear fit $ln(k) = ln(k0) - p_0V - p_1V^2$ for the rest of the nanopores. Error bars indicate standard deviation.

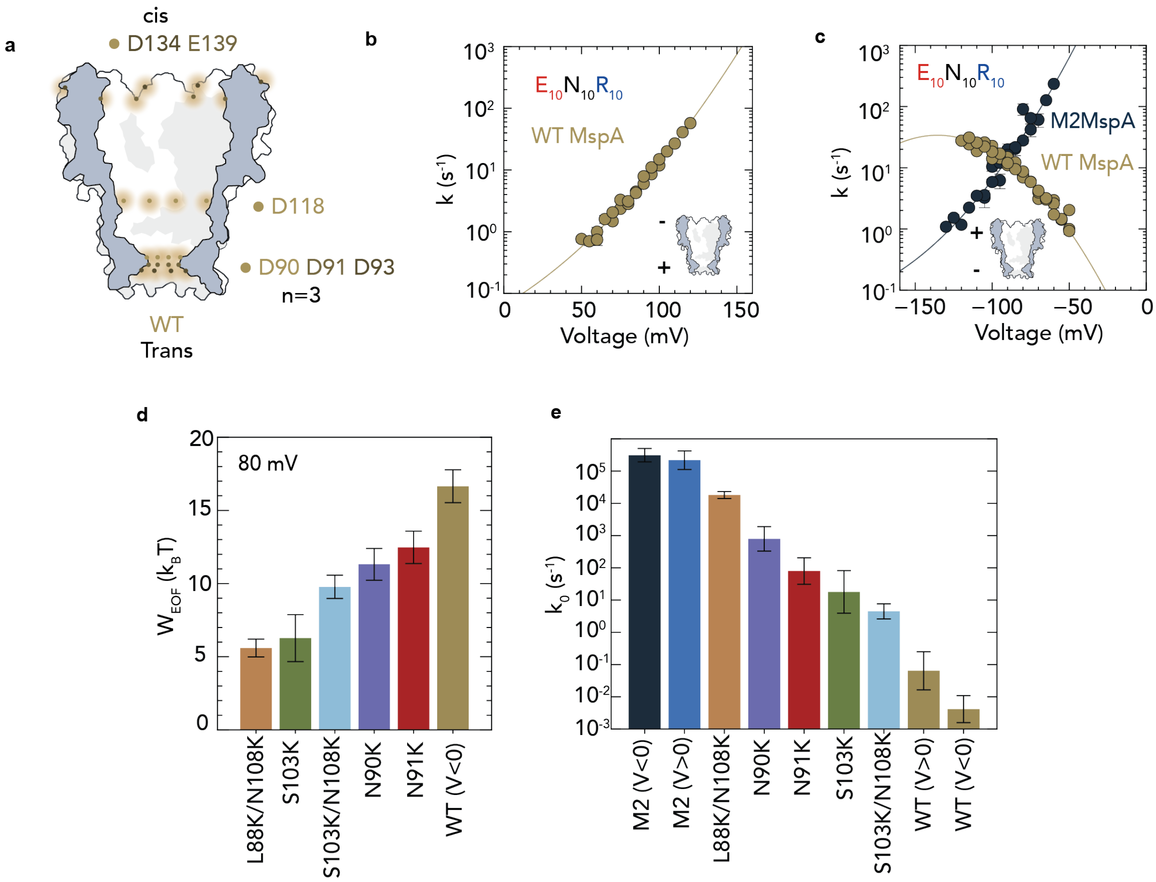


Figure S15: **Work done by EOF measured in the WT MspA nanopore.** a) Positions of the charges in the WT MspA. b) Escape rate vs. voltage measured at positive voltages for the WT MspA. The EOF now runs in the *trans* to *cis* direction, expelling the peptides back into the *cis* reservoir. Error bars indicate standard error of the mean. c) Escape rate vs. voltage measured at negative voltage for the WT MspA and the M2-MspA. In the WT MspA, the EOF runs *cis* to *trans*. For the M2-MspA, the capture rate was low from the *cis* side due to the positive charges at the top of the nanopore that repelled the peptide as it approached with its positive end first. The peptide was instead captured from the *trans* side for the M2-MspA. Both the M2-MspA and the WT MspA was fitted with the equation $ln(k) = ln(k_0) - p_0V - p_1V^2$. Error bars indicate standard error of the mean. d) Bar plot of the work done by EOF for WT MspA and the other variants. Because the M2-MspA at reverse voltage was fitted with a nonlinear term, the equation for the EOF work became: $W_{EOF} = k_BT[(p_{0,M2} - p_{0,mutant})V + (p_{1,M2} - p_{1,mutant})V^2]$. Error bars indicate standard deviation. c) Intrinsic escape rate $k_0$ for the WT MspA and the M2-MspA at negative voltage compared to the other variants. The nonlinear equation $ln(k) = ln(k_0) - p_0V - p_1V^2$ is used to estimate $k_0$ in all pores except M2MspA at positive voltage, which uses the linear equation $ln(k) = ln(k_0) - p_0V$. $k_0$ for the M2-MspA at negative voltage is fairly close to its value at positive voltage. The WT MspA showed the lowest $k_0$ of any pore regardless of voltage. Error bars indicate standard deviation. Fitting parameters are given in Tab.S2 and values for work are given in Tab.S5.

### S3 Supplementary Tables

Table S1: **Fit parameters for** $ln(k) = ln(k_0) - p_0 V$**.** A weighted average were calculated based on the fits from individual replicates. All measurements are for the $E_{10}N_{10}R_{10}$ peptide in 900 mM KCl at positive voltages.

| Nanopore | $\ln(k_0) \pm$ SEM | $p_0 \pm$ SEM ($k_B T$/mV) | $\chi^2_\nu$ | $p$-value |
|---|---|---|---|---|
| M2-MspA | $12.29 \pm 0.38$ | $-0.1146 \pm 0.0040$ | 0.96 | 0.497 |
| L88K/N108K | $9.941 \pm 0.049$ | $-0.04648 \pm 0.00052$ | 1.25 | 0.125 |
| S103K | $3.18 \pm 0.16$ | $-0.0400 \pm 0.0026$ | 2.52 | $1.97 \times 10^{-4}$ |
| S103K/N108K | $3.44 \pm 0.17$ | $-0.0187 \pm 0.0021$ | 4.18 | $3.60 \times 10^{-20}$ |
| N90K | $7.12 \pm 0.16$ | $0.0207 \pm 0.0022$ | 5.54 | $2.25 \times 10^{-26}$ |
| N91K | $4.81 \pm 0.16$ | $0.0359 \pm 0.0025$ | 3.61 | $5.25 \times 10^{-9}$ |

Table S2: **Fit parameters for $ln(k) = ln(k_0) - p_0V - p_1V^2$ for MspA mutants.** A weighted average were calculated based on the fits from individual replicates. All measurements are for the $E_{10}N_{10}R_{10}$ in 900 mM KCl. Voltage is positive where not specified. For pores measured at negative voltages, the voltages were multiplied by -1 before fitting for easier comparison.

| Nanopore | $\ln(k_0) \pm$ SEM | $p_0 \pm$ SEM ($k_BT$/mV) | $p_1 \pm$ SEM ($10^{-4}$ $k_BT$/mV$^2$) | $\chi^2_\nu$ | $p$-value |
|---|---|---|---|---|---|
| M2-MspA | $12.40 \pm 2.25$ | $-0.117 \pm 0.047$ | $-0.125 \pm 2.435$ | 1.00 | $4.48 \times 10^{-1}$ |
| L88K/N108K | $9.80 \pm 0.14$ | $-0.0432 \pm 0.0032$ | $-0.18 \pm 0.17$ | 1.30 | $9.48 \times 10^{-2}$ |
| S103K | $2.88 \pm 0.88$ | $-0.030 \pm 0.029$ | $-0.8 \pm 2.2$ | 1.91 | $1.35 \times 10^{-2}$ |
| S103K/N108K | $1.49 \pm 0.22$ | $0.0367 \pm 0.0060$ | $-3.64 \pm 0.39$ | 1.12 | $2.73 \times 10^{-1}$ |
| N90K | $6.67 \pm 0.44$ | $0.034 \pm 0.012$ | $-0.86 \pm 0.81$ | 4.74 | $6.11 \times 10^{-19}$ |
| N91K | $4.37 \pm 0.55$ | $0.050 \pm 0.018$ | $-1.1 \pm 1.4$ | 3.89 | $4.34 \times 10^{-9}$ |
| M2-MspA ($V \leq 0$) | $12.65 \pm 0.41$ | $-0.1398 \pm 0.0095$ | $3.18 \pm 0.52$ | 4.93 | $2.59 \times 10^{-10}$ |
| WT ($V \leq 0$) | $-5.49 \pm 0.56$ | $0.133 \pm 0.013$ | $-4.91 \pm 0.75$ | 6.56 | $5.23 \times 10^{-24}$ |
| WT ($V \geq 0$) | $-2.75 \pm 0.96$ | $0.034 \pm 0.021$ | $-1.87 \pm 1.11$ | 3.03 | $2.44 \times 10^{-5}$ |

Table S3: **Fit parameters at different salt concentrations.** A weighted average were calculated based on the fits from individual replicates. The M2-Mspa were fit to the linear expression $ln(k) = ln(k_0) - p_0V$ with the exception of the 2 M concentration. The S103K/N108K mutant measurements and the 2 M concentration for the M2-MspA were fit to the nonlinear expression $ln(k) = ln(k_0) - p_0V - p_1V^2$. The peptide $E_{10}N_{10}R_8$ were used in all measurements. All measurements were carried out in the positive voltage range.

| Nanopore | $\ln(k_0/\mathrm{s}^{-1}) \pm$ SEM | $p_0 \pm$ SEM ($k_BT$/mV) | $p_1 \pm$ SEM ($10^{-4}$ $k_BT$/mV$^2$) | $\chi^2_\nu$ | $p$-value |
|---|---|---|---|---|---|
| M2-MspA (500 mM) | $12.8 \pm 1.1$ | $-0.099 \pm 0.011$ | N/A | 2.13 | $1.03 \times 10^{-3}$ |
| M2-MspA (600 mM) | $12.51 \pm 0.25$ | $-0.0917 \pm 0.0027$ | N/A | 2.31 | $1.40 \times 10^{-5}$ |
| M2-MspA (900 mM) | $13.16 \pm 0.16$ | $-0.0976 \pm 0.0015$ | N/A | 1.47 | $4.26 \times 10^{-2}$ |
| M2-MspA (2 M) | $10.27 \pm 0.35$ | $-0.0313 \pm 0.0080$ | $-1.93 \pm 0.45$ | 1.54 | $1.87 \times 10^{-2}$ |
| S103K/N108K (500 mM) | $-0.87 \pm 0.27$ | $0.0427 \pm 0.0077$ | $-2.14 \pm 0.53$ | 1.55 | $1.98 \times 10^{-2}$ |
| S103K/N108K (600 mM) | $0.55 \pm 0.54$ | $0.038 \pm 0.015$ | $-2.04 \pm 0.98$ | 1.32 | $6.72 \times 10^{-2}$ |
| S103K/N108K (900 mM) | $2.28 \pm 0.22$ | $0.0456 \pm 0.0061$ | $-3.14 \pm 0.40$ | 1.71 | $4.84 \times 10^{-3}$ |
| S103K/N108K (2 M) | $7.60 \pm 0.18$ | $-0.0018 \pm 0.0047$ | $-1.71 \pm 0.29$ | 1.72 | $6.67 \times 10^{-3}$ |

Table S4: **Fit parameters for the peptide series $E_{10}N_{10}R_x$ peptides, x = 7, 8, 9, 10.** A weighted average were calculated based on the fits from individual replicates. The linear fit $ln(k) = ln(k_0) - p_0V$ was used for the M2-MspA and the nonlinear fit $ln(k) = ln(k_0) - p_0V - p_1V^2$ was used for the S103K/N108K and the L88K/N108K mutants. All measruements are in 900 mM KCl in the positive voltage range.

| Nanopore | $\ln(k_0) \pm$ SEM | $p_0 \pm$ SEM ($k_BT$/mV) | $p_1 \pm$ SEM ($10^{-4}$ $k_BT/\text{mV}^2$) | $\chi^2_\nu$ | $p$-value |
|---|---|---|---|---|---|
| M2-MspA $E_{10}N_{10}R_{10}$ | $12.29 \pm 0.38$ | $-0.1146 \pm 0.0040$ | N/A | 0.96 | $4.97 \times 10^{-1}$ |
| M2-MspA $E_{10}N_{10}R_9$ | $12.60 \pm 0.11$ | $-0.1063 \pm 0.0012$ | N/A | 1.70 | $9.95 \times 10^{-3}$ |
| M2-MspA $E_{10}N_{10}R_8$ | $13.16 \pm 0.16$ | $-0.0976 \pm 0.0015$ | N/A | 1.47 | $4.26 \times 10^{-2}$ |
| M2-MspA $E_{10}N_{10}R_7$ | $12.85 \pm 0.16$ | $-0.0805 \pm 0.0014$ | N/A | 2.57 | $5.22 \times 10^{-7}$ |
| S103K/N108K $E_{10}N_{10}R_{10}$ | $1.49 \pm 0.22$ | $0.0367 \pm 0.0060$ | $-3.64 \pm 0.39$ | 1.12 | $2.73 \times 10^{-1}$ |
| S103K/N108K $E_{10}N_{10}R_9$ | $1.55 \pm 0.31$ | $0.0451 \pm 0.0087$ | $-3.76 \pm 0.58$ | 0.98 | $5.15 \times 10^{-1}$ |
| S103K/N108K $E_{10}N_{10}R_8$ | $2.28 \pm 0.22$ | $0.0456 \pm 0.0061$ | $-3.14 \pm 0.40$ | 1.71 | $4.84 \times 10^{-3}$ |
| S103K/N108K $E_{10}N_{10}R_7$ | $2.47 \pm 0.16$ | $0.0584 \pm 0.0043$ | $-3.46 \pm 0.27$ | 2.43 | $2.89 \times 10^{-7}$ |
| L88K/N108K $E_{10}N_{10}R_8$ | $9.35 \pm 0.22$ | $-0.0079 \pm 0.0052$ | $-1.09 \pm 0.29$ | 2.23 | $4.32 \times 10^{-6}$ |

Table S5: **Work $W$ done by EOF on the analyte at 80 mV.** $W_{lin}$ is calculated using linear fits for both the reference and the condition through $W_{lin} = k_BT(p_{0,M2} - p_{0,mutant})V$. $W_{curv}$ uses the linear fit for the reference but a nonlinear fit for the measured condition through $W_{EOF} = k_BT[(p_{0,M2} - p_{0,mutant})V - p_{1,mutant}V^2]$, unless marked with * in which case both the reference and the condition uses curved fits through $W_{curv} = k_BT[(p_{0,M2} - p_{0,mutant})V + (p_{1,M2} - p_{1,mutant})V^2]$.

| Nanopore | Reference | Peptide | KCl | $W_{\text{lin}} \pm \sigma_W$ $(k_BT)$ | $W_{\text{curv}} \pm \sigma_W$ $(k_BT)$ |
|---|---|---|---|---|---|
| S103K/N108K | M2-MspA | $E_{10}N_{10}R_{10}$ | 900 mM | $7.67 \pm 0.69$ | $9.78 \pm 0.80$ |
| S103K | M2-MspA | $E_{10}N_{10}R_{10}$ | 900 mM | $5.97 \pm 0.66$ | $6.3 \pm 1.6$ |
| L88K/N108K | M2-MspA | $E_{10}N_{10}R_{10}$ | 900 mM | $5.45 \pm 0.56$ | $5.59 \pm 0.61$ |
| N90K | M2-MspA | $E_{10}N_{10}R_{10}$ | 900 mM | $10.82 \pm 0.65$ | $11.3 \pm 1.1$ |
| N91K | M2-MspA | $E_{10}N_{10}R_{10}$ | 900 mM | $12.04 \pm 0.65$ | $12.5 \pm 1.1$ |
| WT | M2-MspA ($V \leq 0$) | $E_{10}N_{10}R_{10}$ | 900 mM | $10.43 \pm 0.32$ | 16.7*±1.1 |
| S103K/N108K | M2-MspA | $E_{10}N_{10}R_8$ | 500 mM | $8.9 \pm 1.5$ | $10.0 \pm 1.6$ |
| S103K/N108K | M2-MspA | $E_{10}N_{10}R_8$ | 600 mM | $7.90 \pm 0.63$ | $9.1 \pm 1.4$ |
| S103K/N108K | M2-MspA | $E_{10}N_{10}R_8$ | 900 mM | $7.63 \pm 0.37$ | $9.45 \pm 0.47$ |
| S103K/N108K | M2-MspA | $E_{10}N_{10}R_8$ | 2 M | $2.90 \pm 0.14$ | 2.5*±0.7 |
| S103K/N108K | M2-MspA | $E_{10}N_{10}R_{10}$ | 900 mM | $7.67 \pm 0.69$ | $9.78 \pm 0.80$ |
| S103K/N108K | M2-MspA | $E_{10}N_{10}R_9$ | 900 mM | $7.55 \pm 0.36$ | $9.70 \pm 0.60$ |
| S103K/N108K | M2-MspA | $E_{10}N_{10}R_8$ | 900 mM | $7.63 \pm 0.37$ | $9.45 \pm 0.47$ |
| S103K/N108K | M2-MspA | $E_{10}N_{10}R_7$ | 900 mM | $6.68 \pm 0.25$ | $8.89 \pm 0.37$ |
| L88K/N108K | M2-MspA | $E_{10}N_{10}R_8$ | 900 mM | $5.63 \pm 0.27$ | $6.48 \pm 0.51$ |

Table S6: **Ion-permeability values of the different nanopores.** Data were calculated with both the GHK and Nernst equations.[10] The reversal potential was recorded with 2 M KCl in the *cis*-reservoir and 500 mM KCl in the *trans*-reservoir. n $\geq$ 3.

| Nanopore | $V_r \pm \sigma_V$ (mV) | $I_{V=0} \pm \sigma_I$ (pA) | $P_{Cl^-}/P_{K^+}$ $\pm\sigma_P$(GHK) | $P_{Cl^-}/P_{K^+}$ $\pm\sigma_P$ (Nernst) |
|---|---|---|---|---|
| M2-MspA | $-1.65 \pm 0.35$ | $4.35 \pm 0.16$ | $1.12 \pm 0.03$ | $1.11 \pm 0.02$ |
| L88K/N108K | $-6.46 \pm 0.07$ | $14.86 \pm 0.52$ | $1.58 \pm 0.01$ | $1.498 \pm 0.007$ |
| S103K | $-10.43 \pm 0.92$ | $32.64 \pm 1.04$ | $2.12 \pm 0.15$ | $1.94 \pm 0.12$ |
| S103K/N108K | $-12.09 \pm 0.79$ | $39.64 \pm 2.57$ | $2.42 \pm 0.15$ | $2.19 \pm 0.12$ |
| N90K | $-16.64 \pm 1.0$ | $45.21 \pm 1.58$ | $3.57 \pm 0.34$ | $3.11 \pm 0.26$ |
| N91K | $-22.98 \pm 1.00$ | $65.94 \pm 1.18$ | $7.54 \pm 0.02$ | $5.88 \pm 0.73$ |

Table S7: **EOF estimated from the GHK ion-permeability using the equation from Gu *et.al.* 2003.**[4] Estimate is at 900 mM KCl and 80 mV assuming a hydration shell of 6 water molecules.

| Nanopore | $I \pm \sigma_I$ (pA) | $G \pm \sigma_G (mS)$ | $Q_{EOF}$ $\pm\sigma_{Q_{EOF}}$ ($10^9$ molecules $s^{-1}$ ) |
|---|---|---|---|
| M2-MspA | $134.6 \pm 2.0$ | $1.68 \pm 0.02$ | $0.28 \pm 0.07$ |
| L88K/N108K | $171.5 \pm 3.7$ | $2.14 \pm 0.05$ | $1.44 \pm 0.04$ |
| S103K | $133.4 \pm 2.6$ | $1.67 \pm 0.03$ | $1.79 \pm 0.16$ |
| S103K/N108K | $160.9 \pm 0.8$ | $2.01 \pm 0.01$ | $2.50 \pm 0.16$ |
| N90K | $169.0 \pm 1.3$ | $2.11 \pm 0.02$ | $3.56 \pm 0.21$ |
| N91K | $152.2 \pm 1.2$ | $1.90 \pm 0.01$ | $4.36 \pm 0.03$ |

[3] Henrik Bruus. *Theoretical Microfluidics*. Number 18 in Oxford Master Series in Physics. Oxford University Press, Oxford, 2008.

[4] Li-Qun Gu, Stephen Cheley, and Hagan Bayley. Electroosmotic enhancement of the binding of a neutral molecule to a transmembrane pore. *Proceedings of the National Academy of Sciences*, 100(26):15498–15503, 2003.

[5] Dmitriy V Melnikov, Zachery K Hulings, and Maria E Gracheva. Electro-osmotic flow through nanopores in thin and ultrathin membranes. *Physical Review E*, 95(6):063105, 2017.

[6] Michael Rubinstein and Ralph H Colby. *Polymer physics*, volume 23. Oxford university press, New York, 2003.

[7] W. Sung and P. J. Park. Polymer translocation through a pore in a membrane. *Physical review letters*, 77(4):783, 1996.

[8] Chenyu Wen and Shi-Li Zhang. Fundamentals and potentials of solid-state nanopores: a review. *Journal of Physics D: Applied Physics*, 54(2):023001, 2021.

[9] Kherim Willems, Dino Ruić, Florian L. R. Lucas, Ujjal Barman, Niels Verellen, Johan Hofkens, Giovanni Maglia, and Pol Van Dorpe. Accurate modeling of a biological nanopore with an extended continuum framework. *Nanoscale*, 12(32):16775–16795, 2020.

[10] Shouwei Zhang, Jinfeng Wang, Andriy Yaroshchuk, Qiujiao Du, Pengyang Xin, Merlin L Bruening, and Fan Xia. Addressing challenges in ion-selectivity characterization in nanopores. *Journal of the American Chemical Society*, 146(16):11036–11042, 2024.
